# Genetic architecture and cellular basis of flag leaf size variation in barley

**DOI:** 10.1101/2025.04.28.650967

**Authors:** Twinkal Lapasiya, Yanrong Gao, Po-Ya Wu, Amirah Haweit, Delphine van Inghelandt, Benjamin Stich, Asis Shrestha

## Abstract

Flag leaf is the major contributor of photosynthetic assimilates to developing grains. We investigated the genetic architecture and cellular basis of flag leaf length (FLL) and width (FLW) in a multi-parent population comprised of 45 recombinant inbred line (RIL) populations (HvDRR) in barley. Fine mapping of a major quantitative trait locus (QTL) was performed as a first step to isolate the causal gene. Natural variation of FLL and FLW across multi-environment was highly heritable and genotypes adapted to warm climate produced longer and wider flag leaf than those adapted to cooler regions. The variation in flag leaf size was quantitatively inherited and influenced by 24 consensus QTLs of which 17 have not been reported earlier. Further, validation of QTL *qHvDRR-FLS-8* and *qHvDRR-FLS-17* in nearly isogenic RILs demonstrated that these QTLs also controlled length and width of leaves older than flag leaf. The number of epidermal cells primarily determined the differences in FLL and FLW. In addition, we identified the previously unknown effect of genic- and epiallele at *Vrn-H1* on flag leaf size variation in spring barley. Furthermore, we fine-mapped *qHvDRR-FLS-8,* narrowing the interval from 8.7 to 3.5 Mb. In conclusion, our study identified the genomic regions associated with morphological and anatomical variation for leaf size and set the stage to uncover causal genes.

## INTRODUCTION

Photosynthesis is central to plant production and productivity (Long et al., 2006) as it plays a vital role in production and accumulation of organic material in plants (Gautam et al., 2022) and the leaf represents the most important photosynthetic organ (Blum, 1985, Wang et al., 2016). Because of this important role of leaves for photosynthesis, in the 1960s Donald (1968) and Jennings (1964) had proposed a model for enhancing grain yield potential by modifying phenotypic characters such as leaf area (LA) in cereals using “ideotype” breeding. This theory was then supported by the observations that longer flag leaves contributed to higher yields in rice (Rahman et al., 2013) and also, larger and heavier leaf blades led to high grain output in six-rowed barley (Alqudah and Schnurbusch, 2015).

The position of individual leaves affects their contribution to grain yield. During the grain-filling stage of wheat, it was reported that the photosynthetic carbon assimilation of flag leaves (the final leaf to emerge before the spike) supplies >80% of the carbohydrate substrates for starch synthesis (Wu et al., 2012, Fan et al., 2017, Yang et al., 2022). In rice, the flag leaf, produces more than 50% of the carbohydrates that accumulate in grains (Gladun, 1993). A similar observation in barley (Zheng, 1999) suggests that the size of the flag leaf is central to single plant yield. This observation supports the relevance of understanding the genetic architecture of this phenotypic character.

Identification of genes underlying leaf size and shape traits are well advanced in rice. Several genes controlling flag leaf traits have been identified in rice, such as *NARROW LEAF 7 (Nal7)* (Fujino et al., 2008), *NARROW LEAF 1 (Nal1)* (Qi et al., 2008), *SHALLOT-LIKE 1 (Sll1)* (Zhang et al., 2009), *NARROW AND ROLLED LEAF 1 (Nrl1)* (Hu et al., 2010), *SEMI-ROLLED LEAF 1 (Srl1)* (Xiang et al., 2012, Li et al., 2017). By mutant analysis of these genes, it has been found that changes in leaf shape and size resulted from changes in cell division and/or expansion. For instance, Jiang et al. (2015) have found that leaf length and width are reduced by 50% in *nal1-2* and *nal1-3* rice mutants which was caused by suppression of cell division. In addition, natural variation for flag leaf area and related traits was studied using quantitative trait locus (QTL) mapping in rice. Such studies identified additional genomic regions compared to those from mutant based analyses suggesting that the genetic network underlying leaf size variation in rice is not yet fully uncovered (Zhang et al., 2015, Wu et al., 2017, Tang et al., 2018, Wen et al., 2020, Wang et al., 2022, Wang et al., 2023).

In comparison to the large number of leaf mutants available in rice and maize (Neuffer et al., 1997) only a few leaf mutants are available for barley and even fewer have been characterized. The available barley leaf mutants have been categorized as having narrow (e.g., *angustifolium*, *fol*), wide (e.g., *Broad leaf1*, *Blf1; Broad leaf13, Blf13*), long (e.g., *Curly3*, C*ur3*), or short leaves (e.g., *Curly dwarf1*, *Cud1*; *Elongation1-5*, *Elo1-5*). For some of these mutants, chromosomal positions have been determined (Druka et al., 2011).

Yet, the underlying genes for only three barley leaf mutations have been identified and functionally characterized. The recessively inherited *narrow leafed dwarf1* (*nld1*) mutant is characterized by reduced plant height and narrow leaf blade with reduced leaf width and length (Yoshikawa et al., 2016). The narrow-leaf phenotype is caused by a reduction of the number of cells across the lamina of all leaves. This observation indicates that normal *Nld1* function is required to promote medial-lateral, but not proximal-distal, lamina growth throughout plant development (Yoshikawa et al., 2016). Positional cloning uncovered that the *Nld1* gene encodes a WUSCHEL-RELATED HOMEOBOX (WOX) transcription factor, related to the maize proteins *NARROW SHEATH1* (*Ns1*) and *Ns2* (Yoshikawa et al., 2016). In contrast to *Nld1*, the barley mutants *broad leaf1* (*Blf1*) and *broad leaf13 (Blf13)* are characterized by wide leaf blades compared to the wild-type which is caused by an increased number of cells along the medial-lateral axis (Jöst et al., 2016, 2024). *Blf1* has no significant effect on the width of the leaf sheath. In contrast, the width of the palea and lemma of *blf1* mutants increases, which suggests the existence of shared genetic mechanisms to control the medial-lateral growth between these organs and leaf blade. The effect of *blf1* on blade width appears from plastochron 6 (P6) onward, indicating that *Blf1* functions to limit cell proliferation in the medial-lateral axis during blade outgrowth, but does not affect the recruitment of leaf founder cells as *Ns1/2* does (Jöst et al., 2016, 2024). The *Blf1* locus encodes an INDETERMINATE-domain protein that is expressed in the nuclei of shoot apical meristem, epidermal, and sub-epidermal cells at the base of P2 and P3 leaf primordia and later throughout the epidermis (P5/P6), especially in correspondence with presumptive veins (Jöst et al., 2016). The T189P substitution in the conserved motif of *HvHNT1* protein caused that *Blf13* mutants produce wider leaf primordia (after passing P2 stage) and hence, wider leaf blades than wild type (Jost et al., 2024). To our knowledge, no earlier studies have explored the presence of natural allelic variation at *Nld1*, *Blf1, and Blf13*.

In addition to these three mutant-based approaches mentioned above, several studies have employed the existing natural variation for leaf size in cultivated barley to unravel its genetics using genome-wide association mapping. Digel et al. (2016) have assessed the leaf blade width and length variation in a diverse panel of European winter barley cultivars and observed that the *Ppd-H1* locus is associated with the studied traits. The recessive late flowering allele at *Ppd-H1* correlates with larger blade width and length. A similar observation has been reported by Alqudah et al. (2018) in a spring barley association panel.

In addition, some studies on QTL for flag leaf length and width have been conducted in bi-parental populations (Gyenis et al., 2007, Xue et al., 2008, Liu et al., 2015, Vafadar Shamasbi et al., 2017, Niu et al., 2022). The before mentioned studies detected various QTLs for flag leaf size across all the chromosomes of barley. Some studies (Liu et al., 2015; Niu et al., 2022) detected QTLs which co-localized with those of the other studies (Gyenis et al., 2007, Xue et al., 2008, Du et al., 2019). Furthermore, many QTLs have been detected around *Vrs1* locus (Liu et al., 2015). One of the reasons of the relatively few detected novel QTLs in the above mentioned studies might be because of the limited genetic variation captured by a single bi-parental population.

The objectives of this study were to (i) assess the genetic complexity of flag leaf size variation using a multi-parent population (MPP) resource, the double round-robin population of barley (HvDRR), (ii) identify the allelic series at each QTL and candidate genes for the detected QTLs by local association mapping exploiting genome-wide variant information of parental inbreds as well as classical fine mapping, and (iii) assess for a subset of the QTL the cellular cause of flag leaf size variation as well as their pleiotropic effects on plant stature.

## Material and Methods

### Plant materials

Our study was based on the spring barley diversity panel described by Haseneyer et al. (2010). In addition, we have selected from this panel the 23 inbreds that maximize a genotypic and trait diversity index (Weisweiler et al., 2019) and crossed them in the double round-robin design (Stich, 2009). From each of the 45 crosses, a RIL population was developed following the single seed descent principle. The 50K barley single nucleotide polymorphism (SNP) array (Bayer et al., 2017) was used to genotype 35–146 RILs from each of the 45 HvDRR sub-populations in the F4 generation. As described by Casale et al. (2022), the genetic map for individual HvDRR sub-populations, and consensus map across the HvDRR population was established, and used for linkage mapping described below.

### Field experiment and data collection

From 2019 to 2021, each RIL was grown in field experiments at two different locations in Germany: Köln and Mechernich in 2019 and 2020; and Düsseldorf and Mechernich in 2021. We followed an augmented incomplete block design with one replicate for each RIL in each of the six environments as well as 16-20 replicates for each of the 23 parental inbreds. The RILs from a single HvDRR sub-population were sown in a randomized way in adjacent rows or columns with 30 seeds per entry in a 160 cm long row. The spring barley diversity panel with 224 inbreds was included in each environment.

The flag leaf length (FLL) of the main culm in cm was collected from three to five random plants of each entry in each of the six environments. In the years 2020 and 2021, from the same leaf, also the flag leaf width (FLW) was assessed in cm.

### Statistical analyses

Data was processed and statistical analyses were performed using R software (R Core Team 2023). Upon the removal of outliers with residuals larger than 2.5 times the standard deviation of the residuals from a fitted linear model (equation **1**), the measurements made for FLL and FLW in the above described experiment were analyzed using the following mixed model:

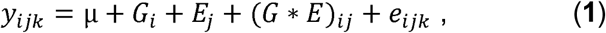

where, *y*_*ijk*_ was the observed phenotypic value for the *i*^*th*^ genotype in the *j*^*th*^ environment for the *k*^*th*^ replication, μ the general mean, *G_i_* the effect of the *i*^*th*^ genotype, *E_j_* the effect of the *j*^*th*^ environment, (G * E)*_ij_* the interaction of *i*^*th*^ genotype with *j*^*th*^ environment, and *e*_*ijk*_ the residual. For the calculation of adjusted entry means, the genotype effect *G* was considered as fixed and the other effects were considered as random. For the calculation of the broad sense heritability on an entry mean basis (*H^2^*), the genotypic variance σ^*2*^_*g*_ was calculated based on the above model, but with a random genotype effect. *H^2^* was then calculated according to Piepho and Moöhring (2007).

Further, we performed a single population QTL analysis for FLL and FLW in the 45 HvDRR sub-populations using the R/qtl package (Broman et al. 2003) as described in detail by Shrestha et al. (2022). First, genotype probabilities of every 1 cM were estimated, followed by a genome-wide scan for QTL using Haley-Knott regression. A forward search algorithm was then applied to perform multiple QTL mapping. A permutation test with 4000 runs was performed, and the logarithm of the odds (LOD) score at a 0.05 significance level was set as a threshold for QTL detection. QTL were added to the model as long as these were significant at the before mentioned threshold.

A 1.5 LOD drop sets the confidence interval on either side of the detected QTL (Dupuis and Siegmund, 1999). The co-localized QTLs for FLL and FLW with overlapping confidence interval detected across the HvDRR sub-populations were summarized as consensus QTLs. The confidence interval of consensus QTLs was defined across sub-populations and phenotypes by using the shortest possible confidence interval of co-located QTLs and referred as flag leaf size (FLS) associated loci detected in HvDRR population (*qHvDRR-FLS*). The percent of explained phenotypic variance (PEV) for these consensus QTLs was summarized as the range of the lowest and highest PEV explained by the individual populations belonging to the respective consensus QTLs.

We also performed multi-parent population (MPP) QTL analysis (Garin et al., 2017) using a consensus map across the RILs of 45 HvDRR sub-population. Parental model was applied to estimate the multi-QTL effect, with the assumption that (a) each parent contributes a unique allele at QTL, and (b) the effect of parental inbred is constant across the sub-populations involving the parent in question. QTL scanning for FLL and FLW was performed according to the procedure described in Shrestha et al. (2022).

### Genomic prediction

Genome-wide prediction for FLL and FLW was performed by genomic best linear unbiased prediction (GBLUP) using the following model (VanRaden, 2008):

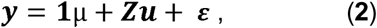

where ***y*** was the vector of the adjusted entry means of the HvDRR population for FLL and FLW, **1** the unit vector, ***µ*** the general mean, **Z** the incidence matrix of genotypic effects, ***u*** the vector of genotypic effects that were assumed to be normally distributed with 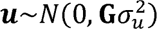, where G denotes the relationship matrix calculated on the basis of the SNP marker data, and 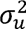 the genetic variance, and E the vector of residuals following the normally distribution 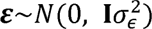, where I was the identity matrix and 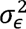 the residual variance. In this study, only additive effects were considered. GBLUP model was performed using R package sommer (Covarrubias-Pazaran, 2016). Five-fold cross validation was applied to evaluate the performance of the GBLUP model. Prediction ability was estimated as the Person’s correlation coefficient between the observed and genomic estimated breeding value (GEBV). The median prediction ability across the five-fold cross validation was calculated for each of the 50 replicates. The percentage of phenotypic variance explained by the GBLUP model was the square of the median prediction ability.

### Candidate gene analysis

We carried out candidate gene analysis for the consensus QTLs that were smaller than 10 Mb. For these QTLs, the list of high confidence genes was obtained from the plant genomics and phenomics research data repository from IPK Gatersleben, Germany. (https://edal-pgp.ipk-gatersleben.de/). The annotation and coordinates were based on the Morex v3 gene models (Mascher et al., 2021). The genes that displayed polymorphisms among the concerned parental inbreds were then selected using the variant calling data gathered from the whole genome sequencing project of the 23 parental inbreds (Weisweiler et al., 2022) and were then designated candidate genes. Further, we looked for non-synonymous (amino acid substitutions) sequence variants categorized as tolerated or deleterious mutations; indels in the coding region; and indels and structural variants (SVs) in the 5’-end regulatory region (5 kb upstream of the start codon).

### Detailed characterization of the effect of QTLs *qHvDRR-FLS-8* and *qHvDRR-FLS-17* on leaf and cell properties

We validated two QTLs, namely *qHvDRR-FLS-8* and *qHvDRR-FLS-17* and dissected the epidermal cell size and number on the leaf blade. Four RILs, two carrying Georgie allele (Hv.2018S.7.02932, Hv.2018S.7.02664) and two carrying HOR8160 allele (Hv.2018S.8.02731 and Hv.2018S.7.02921) at *qHvDRR-FLS-8* were evaluated. Georgie and HOR8160 are the parental inbreds of the HvDRR14 sub-population. At *qHvDRR-LS-17*, three RILs, namely Hv.2018S.7.03209, Hv.2018S.7.03211 (Georgie allele) and Hv.2018S.7.03369 (Lakhan allele) from HvDRR33 sub-population were used for QTL validation. The pair of RIL for a QTL was selected such that they were polymorphic at the QTL of interest but monomorphic at all other QTLs detected in that sub-population and furthermore was monomorphic for the highest number of molecular markers for the rest of the genome. Six plants of each of the seven RILs and three parental inbreds were grown in a lattice design under controlled conditions. Each plant was grown in a 1.5 litre pot filled with a mixture of sand and soil (30:70 w/w peat-based soil “Einheitserde Deklarationstyp 3”). The day length was set to sixteen hours followed by eight hours darkness. The light intensity was set to 300 μmol m^−2^ s^−1^, 50 cm away from the light source and the relative humidity to 55%. The temperature under daylight was 20°C and the night temperature was 18°C.

We measured the length and width of the flag leaf (FL), second (L2), third (L3), and fourth (L4), fully expanded leaf from the top of the main culm. The collected data of leaf size were then analyzed using the following fixed model:

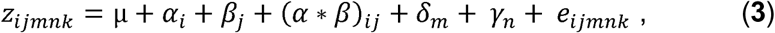

where, *z_ijmnk_* was the observed phenotypic value for *i*^*th*^ genotype for the *j*^*th*^ leaf type at the *m*^*th*^ row and the *n*^*th*^ tray for the *k*^*th*^ replication, μ the general mean, *α_i_* the effect of the *i*^*th*^ genotype, *β_j_* the effect of the *j*^*th*^ leaf type, (a * *β*)*_ij_* the interaction of *i*^*th*^ genotype with *j*^*th*^ leaf type, *δ_m_* the effect of *m*^*th*^ row, *γ_n_* the effect of *n*^*th*^ tray, and *e_ijmnk_* the residual. For the calculation of adjusted entry means, all the effects were considered as fixed.

From the same plants, leaf imprints were taken from the top three leaves for QTL *qHvDRR-FLS-8* and top two leaves for QTL *qHvDRR-FLS-17* to assess cell size and cell number. We prepared epidermal imprints from the middle of the adaxial side of the leaf in direction of the leaf length. This was realized by applying a film of transparent nail polish which was then transferred with transparent adhesive tape to microscope slides. Images of the epidermal cells from the imprints were recorded from two field of views using a stereo microscope Nikon SMZ18 equipped with a Nikon DS-Fi2 camera at 10X magnification. The software ImageJ (Schneider et al., 2012) was used to assess the length of lateral cells and count the number of cell files. The length was measured for all the lateral cells per row per imprint bordering the stomata cell files (Fig. S1). The collected cell dimensions were then analyzed using model (**3**).

### Fine mapping of QTL *qHvDRR-FLS-8* on chr2H

We developed a high-resolution mapping population by crossing two RILs from the HvDRR14 sub-population that were also included in the above described experiment, namely Hv.2018S.7.02932 and Hv.2018S.7.02921 that harbored Georgie and HOR8160 allele at QTL *qHvDRR-FLS-8* on chromosome 2H. Then, we genotyped the left border (LB) and the right border (RB) of the QTL interval in 1890 F2 progenies using Kompetitive allele specific PCR (KASP) markers (Table. S1). DNA was extracted from 7-10 days old seedlings for genotyping, and *KASP*^TM^ assay was performed using Quantstudio5 real-time PCR machine (Thermo Fischer Scientific). The recombinants between LB and RB were transplanted to 1.5 litre pots. The recombinants were further genotyped inside the QTL interval (Table S1) and were grown in the greenhouse until maturity.

Further, the F3 progenies derived from the above mentioned recombinants were grown in the field between March to July 2024 at Groß Lüsewitz in Germany, following an augmented block design with two replicates. The experiment contained two blocks (the replicates) with 30 seeds per recombinant in a 160 cm long row in each block, each recombinant appeared only once in one block. At heading stage, when the flag leaf was fully developed, leaf length was evaluated from three different randomly selected plants per row.

Then, phenotypic data of F3 progenies were analyzed using the following model:

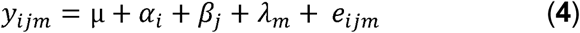

where, *y_ijm_* was the observed phenotypic value for *i*^*th*^ genotype in the *j*^*th*^ block at the *m*^*th*^ range, μ the general mean, *α_i_* the effect of the *i*^*th*^ genotype, *β_j_* the effect of the *j*^*th*^ block, *λ*_*m*_ is the effect of the *m*^*th*^ range and *e_ijm_* the residual. For the calculation of adjusted entry means, all the effects were considered as fixed. We performed marker trait correlation with the adjusted entry means of FLL and the genotypic data of the F2 progenies using an additive and dominance effects model.

## Results

### Phenotypic variation for flag leaf length and width

Flag leaf length and width phenotypes were collected across six and four environments, respectively, for about 3870 RILs from the 45 HvDRR sub-populations as well as the diversity panel. The parental inbreds were replicated between 16 to 20 times in each environment as repeated checks to account for field heterogeneity. We observed a significant variation in the phenotype FLL and FLW among the inbreds of the diversity panel originating from different continents, with genotypes from South Asia and East Asia producing comparatively longer and wider flag leaves (Fig. 1). Smaller flag leaves were common among the genotypes from Europe except southern Europe. Notably, we also observed a considerable diversity for FLL and FLW across the RILs of the HvDRR population. The adjusted entry means of RILs across six environments for FLL varied between 5.6 and 19.1 cm (Fig. 2). For FLW, the range on a proportional scale was even larger with 0.25 to 2 cm (Fig. 2). The range of FLL and FLW of RILs from 45 sub-populations was the same as those observed for the diversity panel. The sub-population with the highest diversity for FLL was HvDRR43 with a coefficient of variation (CV) of 18.8%, whereas HvDRR08 had the lowest CV (7.4%). The coefficient of variation of FLL for the diversity panel (14%) fell between the above mentioned two extreme cases. This trend was different for FLW, where the highest CV was observed for the diversity panel with 41.6%. The coefficients of variation for the HvDRR sub-populations were lower ranging from 12.8 (HvDRR07) to 28% (HvDRR47) (Table S2). The majority of these differences were due to genetics as the broad sense heritability on an entry mean basis was 0.77 for FLL and 0.58 for FLW (Table 1). We observed a correlation coefficient of 0.62 (Fig. S2a) for the correlation between FLL and FLW across all the evaluated entries. However, the level of association differed considerably between the different characterized material groups. The correlation coefficient was 0.62 for the diversity panels and ranged from -0.24 to 0.87 for the individual HvDRR sub-populations (Fig. S2a, b and Fig. S3). In 40 HvDRR sub-populations and the diversity panel, the correlation between FLL and FLW was significantly (P < 0.05) different from 0 (Fig. S3).

**Fig. 1.**
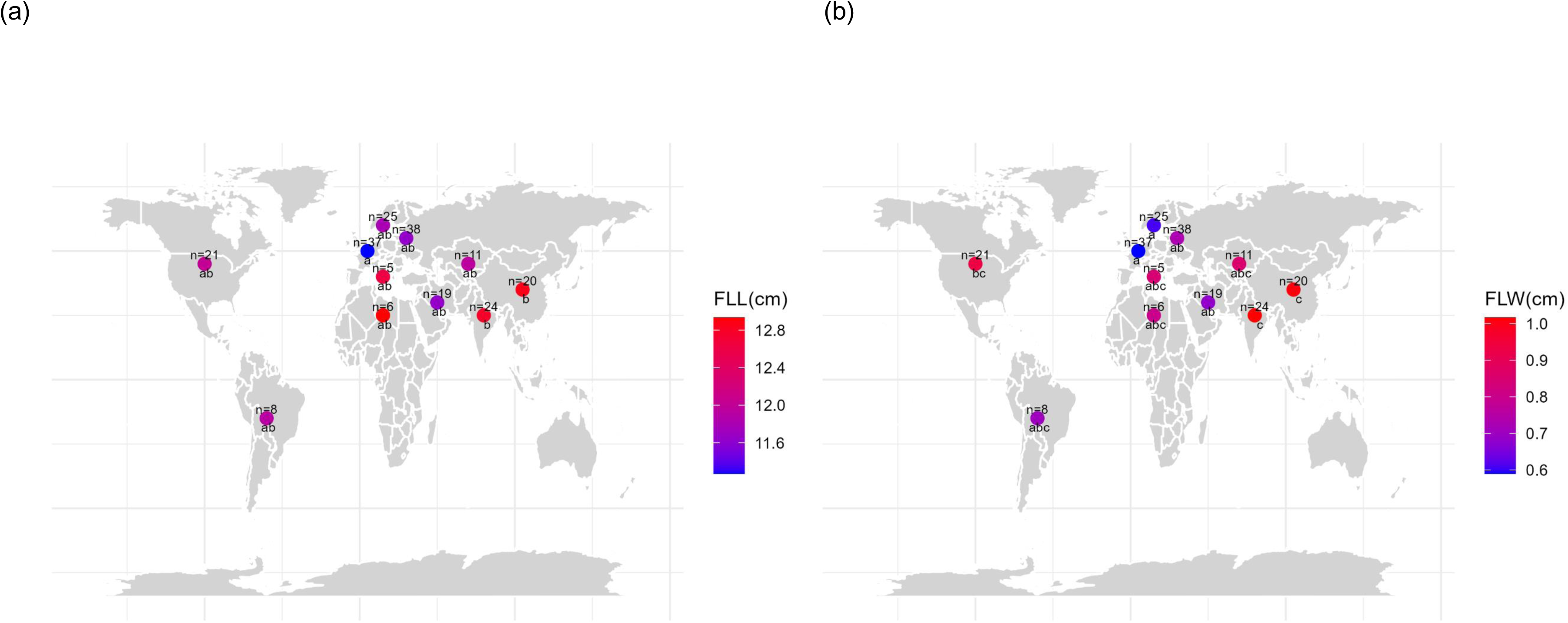
Variation on flag leaf size based on the geographical origin. Heat map of adjusted entry means for (a) Flag leaf length (FLL) in cm and (b) Flag leaf width (FLW) in cm in barley diversity panel according to their geographical origin. Indexed letters below the dots indicate significant differences in the trait value among the accessions from respective geographical region (*p* < 0.05) not sharing the same letter. The values above the dots denote the number of accessions from the diversity panel belonging to the respective geographical region.

**Fig. 2.**
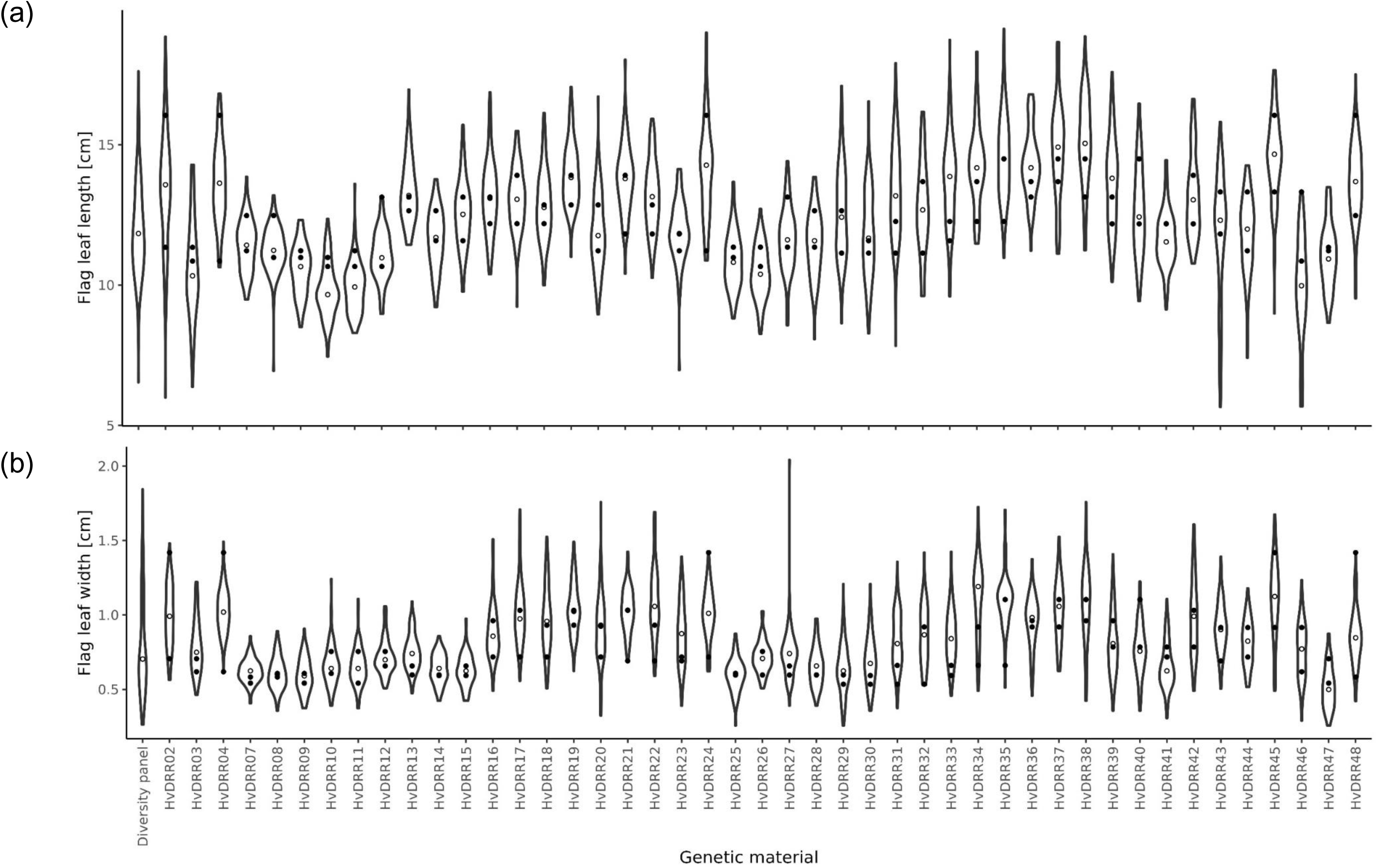
Violin plot for flag leaf size across 45 HvDRR sub-populations. (a) Flag leaf length (FLL) (b) flag leaf width (FLW) adjusted entry mean within each sub-population, with wider regions indicating higher density. The white dot within each violin denotes the median values across all RILs and the two black dots in each violin highlight the adjusted entry means of FLL and FLW of the parental inbreds of each sub-population.

**Table 1:** Variance components and broad-sense heritability (*H^2^*) of flag leaf length and flag leaf width.

| Phenotype | Genotype | Environment | G*E | Error | $H^2$ |
| --- | --- | --- | --- | --- | --- |
| Flag leaf length | 2.82 | 1.18 | 0.76 | 3.62 | 0.77 |
| Flag leaf width | 0.034 | 0.01 | 0.01 | 0.04 | 0.58 |

### QTLs associated with flag leaf size

First, we performed MPP analyses that detected 15 and 12 QTLs associated with FLL and FLW, respectively. Seven of these QTLs were associated with both FLL and FLW (Fig. 3). In the next step, we performed sub-population specific QTL analysis and detected a total of 79 QTLs for FLL and FLW. The QTLs were detected in 29 and 15 of the HvDRR sub-populations for FLL and FLW, respectively (Fig. 4). Among the detected QTLs, 36, 11, and 12 were located on chromosomes 2H, 3H, and 5H, respectively. The number of QTLs on the remaining chromosomes varied between 4 and 6 (Fig. 4, Table S3, S4). Except for a locus on chr2H (*qFL2-2*, *qFW2-2*), all the MPP QTLs were detected at the same position in the sub-population specific analyses which illustrated the accuracy of the QTLs identified in the study. The percentage of explained variance (PEV) for individual QTLs varied from 3 to 46% (Table S3, S4). About 77% of the QTLs detected in our study explained ≥ 10% of the phenotypic variance for the associated characteristics (Table S3, S4). We further summarized these 79 QTLs into 24 consensus QTLs based on their overlapping confidence intervals. Of these 24 consensus QTLs, 12 QTLs were shared between FLL and FLW. Moreover, 10 QTLs were only associated with FLL of which nine were unique to individual sub-populations. The remaining two consensus QTLs were detected only for FLW of which one QTL was unique to sub-population HvDRR31 (Table S5). The percentage of phenotypic variance explained by a GBLUP model across all genome-wide SNPs across the entire HvDRR population was 61.2% (FLL) and 51.9% (FLW).

**Fig. 3.**
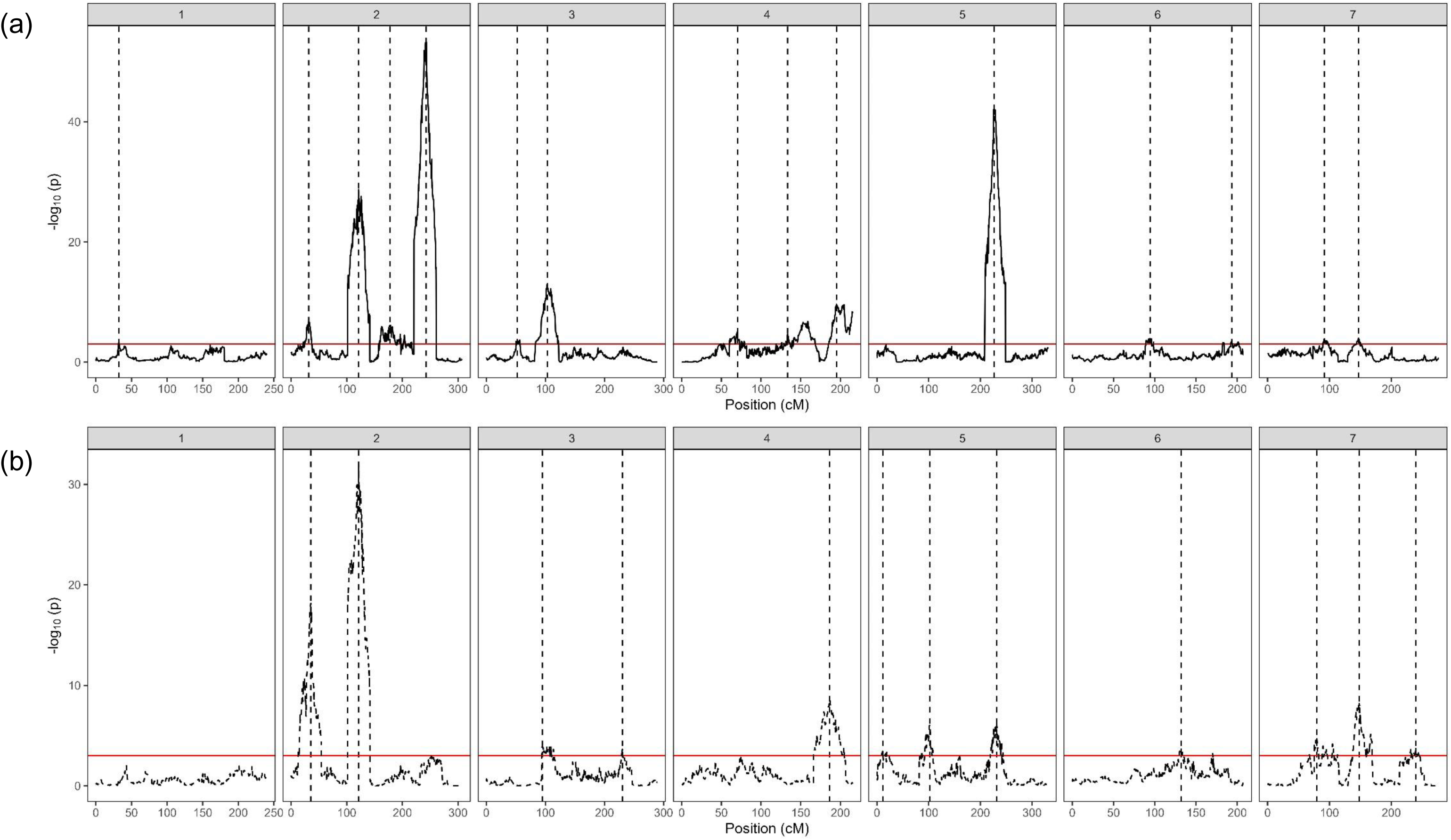
Profiles of quantitative trait loci (QTLs) associated with flag leaf size from multi-parent population analysis across the 45 HvDRR sub-populations using the parental model. QTL profile for (a) flag leaf length, and (b) flag leaf width. The QTLs were detected across all seven chromosomes using composite interval mapping [significant threshold of –log_10_(p)=3]. The vertical dashed lines indicate the positions of the detected QTLs.

**Fig. 4.**
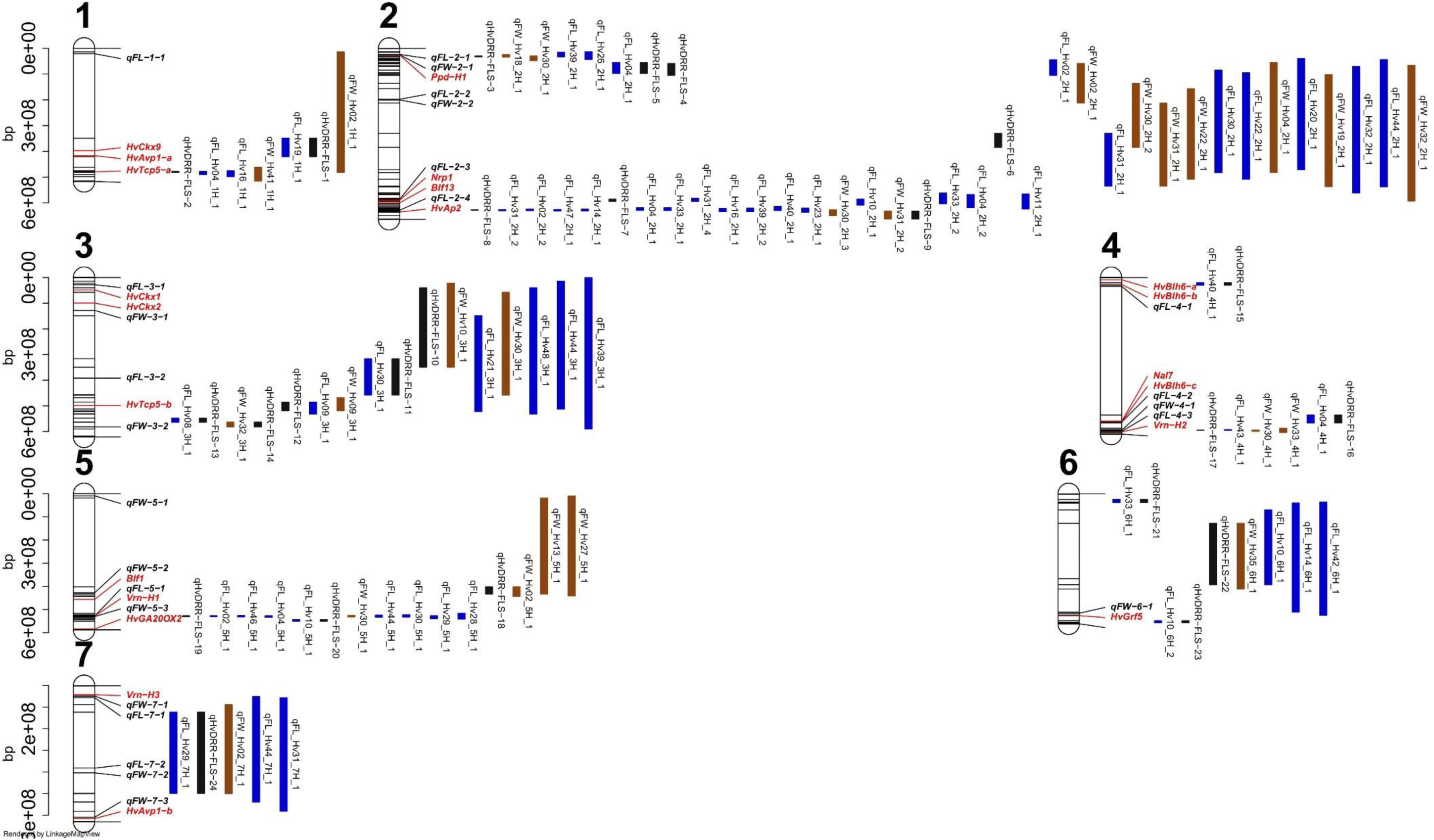
Confidence interval of QTLs detected in individual HvDRR sub-populations. Flag leaf length (FLL) was assessed in six environments and flag leaf width (FLW) in four environments. The blue, brown and black color bar indicates the physical interval of QTLs associated with FLL, FLW and consensus QTLs, respectively. The positions of known genes controlling developmental phenotypes such as leaf size, flowering time, plant height, and spikelet branching are indicated in italic and red font. The positions of QTLs detected from multi-parent population analysis using parental model are indicated in italic and black font. The physical positions of the top and bottom five markers of the genetic map, and the markers flanking the QTL interval are indicated in the plot as black line in the sketched chromosomes.

Next, we estimated the additive effect of each parental inbreds across the MPP QTLs. The QTL allele effects ranged from -0.66 to 0.5 cm and -0.09 to 0.08 cm for FLL and FLW, respectively (Fig. S4). In addition, the crossing design of HvDRR population allows to estimate the allelic series for QTL effect discussed above. At each MPP QTL, we observed an allelic series for both FLL and FLW (Fig. 5). This observation illustrated that the phenotypic variation for FLL and FLW in HvDRR population was controlled by the cumulative effect of minor-effect alleles and a few large effect alleles (Fig. 5 and Fig. S4).

**Fig. 5.**
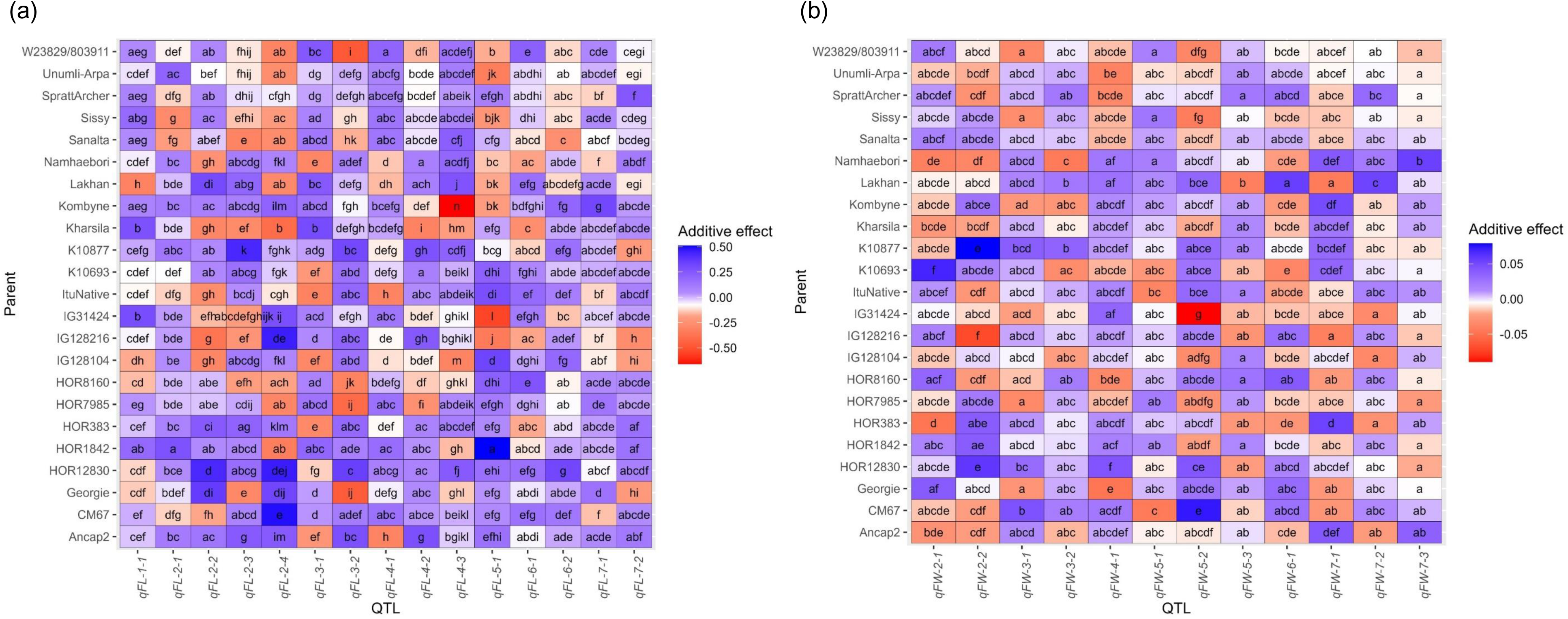
Heat map of the effects of the parental inbreds at the quantitative trait loci (QTLs) detected through multi-parent population analysis using a parental model. Multiple comparisons of the standardized allele effect for (a) flag leaf length (FLL) and (b) width (FLW) at corresponding QTLs. The standardized allele effect for an inbred is the difference between the mean of the estimated allele effect for 23 inbreds and the estimated additive effect of the corresponding inbred. The color code indicates the magnitude of the standardized allele effect. Indexed letters indicate the significant difference (p ≤ 0 05) between the genotypes not sharing the same letter by Tukey’s HSD test.

### Candidate genes associated with flag leaf length and width

The variant calling data from the whole-genome sequencing of the 23 parental inbreds, including SNP annotations, indels, and predicted SVs (Weisweiler et al., 2022) were used to select the candidate genes in the consensus QTL interval that were smaller than 10 Mb, which was the case for nine QTLs (*qHvDRR-FLS-2, qHvDRR-FLS-3, qHvDRR-FLS-7, qHvDRR-FLS-8, qHvDRR-FLS-15, qHvDRR-FLS-17, qHvDRR-FLS-19, qHvDRR-FLS-20* and *qHvDRR-FLS-23)*. For these nine consensus QTLs, we found a total of 900 positional candidate genes. The number of positional candidate genes for each consensus QTLs varied between 11 and 206 (Table S6).

Out of the 24 consensus QTLs, 12 contained the closest barley orthologues of genes involved in developmental processes in barley or other crops (Table S5, S7). Allelic variations in the regulatory or coding region for some of these orthologues were found among the parental inbreds of QTL bearing sub-populations (Table S5, S6). For example, we found InDels in the promoter region of *Tcp5 and Grf5* gene residing in *qHvDRR-FLS-2* which is known to be involved in defining organ size in Arabidopsis (van Es et al., 2018, Yu et al., 2021, Horiguchi et al., 2005). At *qHvDRR-FLS-2,* we found 5 bp to 13 bp deletions in the promoter region of *Tcp5* gene in HOR1842, HOR12830, K10693 (contributing to longer/wider FL) compared to ItuNative (contributing to shorter/narrower FL) indicating possible cis-regulatory effect (Fig. S5). Likewise, 6 bp deletions in the 5’ UTR region and 25 bp deletions in the first intron of CM67 (contributing to longer FL) compared to ItuNative (contributing shorter FL) in *Grf5* was the promising candidate gene for *qHvDRR-FLS-15* (Fig. S6). In addition, we also detected QTLs for FLL and FLW at major flowering time loci in barley (Fig. S7, Tables 2, S8 and S9). The hyper methylated epiallele and intron variation allele *Vrn-H1-3* associated with earlier flowering contributed to longer FL (Table 2, Fig. S7).

**Table 2:**
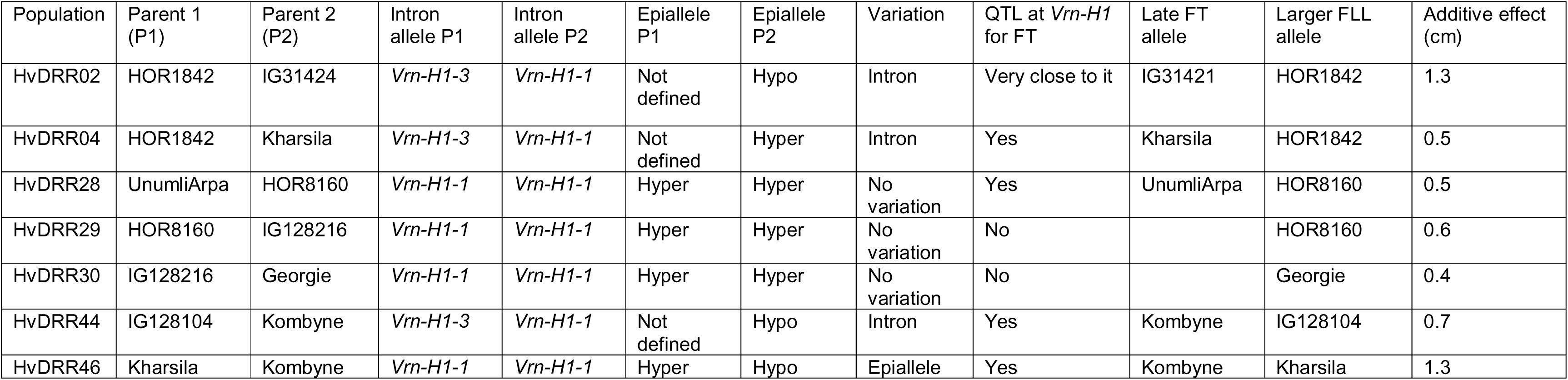
The allele of the *Vrn-H1* gene among the parental inbreds. The table summarizes the deletion and differential methylation (epiallele) for the parental inbreds of those sub-populations where a QTL was detected for flag leaf length (FLL) and width (FLW) at the *Vrn-H1* locus. The nomenclature for the intron deletion detected was adapted from Hemming et al. (2008) and the epiallele was adapted from Kühl et al. (2024). The flowering time QTLs detected by Cosenza et al. (2024) were used to compare the association of the effect of late or early *Vrn-H1* flowering time (FT) allele with FLL.

### Validation and characterization of QTL *qHvDRR-FLS-8* and *qHvDRR-FLS-17* under controlled conditions

*qHvDRR-FLS-8* on chr2H was one of the major effect QTLs for FLS with PEV ranging from 8-32% across the individual HvDRR sub-populations (Table S5). We validated the effect of *qHvDRR-FLS-8* on FLL in the HvDRR14 sub-population by comparing RILs Hv.2018S.7.02932 and Hv.2018S.7.02664 carrying Georgie allele with RILs Hv.2018S.8.02731 and Hv.2018S.7.02921 carrying HOR8160 allele at *qHvDRR-FLS-8* while the rest of the genome was close to isogenic. We observed that the FL, L2, and the L3 of the RILs with Georgie allele were significantly longer (P< 0.05) compared to the RILs carrying the HOR8160 allele at this QTL. This trend was in accordance with the QTL effect observed in the QTL analysis but in contrast to the length of the leaves in the parental inbreds, where Georgie had shorter leaves than HOR8160 (Fig. 6a-c). The difference in the parental effect is due to that several QTL with varying direction of the QTL effects segregate among them whereas only one QTL segregates among the individuals of our validation experiment.

**Fig. 6.**
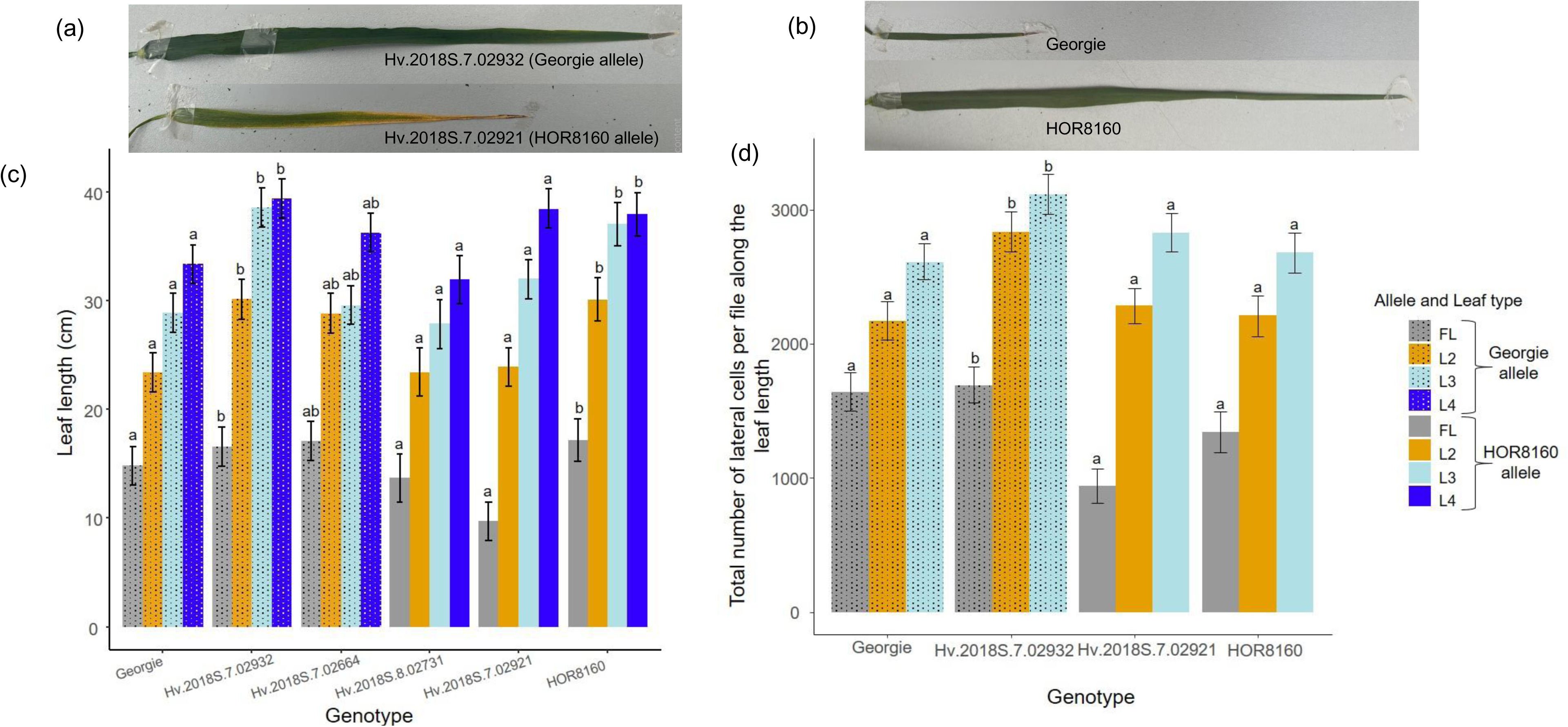
Validation of a major QTL *qHvDRR-FLS-8* associated with flag leaf length. *qHvDRR-FLS-8* was detected on chr2H and the validation was done in the HvDRR14 sub-population. Images of flag leaf of (a) Recombinant inbred lines (RILs) and (b) Parental inbreds for visual comparison. (c) The length of flag leaf and older leaves in RILs and parental inbreds of sub-population HvDRR14. RILs were polymorphic for *qHvDRR-FLS-8* but isogenic for the other QTLs detected in the HvDRR14 sub-population. (d) The number of lateral cells per file along the leaf length. Bar indicates leaf length/total number of lateral cells per file along the leaf length (AEM ±SE, n = 6). Grey, yellow, light blue, and dark blue bar represents flag leaf (FL), second (L2), third (L3), and fourth leaf from the top (L4), respectively. Patterned bars in the graph represent Georgie or RILs with Georgie allele and non-patterned bars represent HOR8160 or RILs with HOR8160 allele at *qHvDRR-FLS-8*. Bars of the same color not sharing the same letter above the bar are significantly different from each other (*p* < 0.05). AEM, adjusted entry mean; SE, standard error.

Next, we measured the length of lateral cells in the *qHvDRR-FLS-8* RIL pairs and the parental inbreds and further used this data to estimate the number of lateral cells per file across the leaf length. This was done in order to determine if leaf length differences are due to longer lateral cells or more lateral cells. We observed a significant difference in the length of the lateral cells between the parental inbreds (Fig. S8). However, the RIL pairs with contrasting allele at *qHvDRR-FLS-8* showed a significant difference in the number of lateral cells per file across leaf length which indicates that at this QTL the difference in the leaf length is due to difference in cell proliferation (Fig. 6d).

*qHvDRR-FLS-17* on chr4H was another major effect QTL for FLS with PEV ranging from 13-28% across the HvDRR sub-populations (Table S5). We validated *qHvDRR-FLS-17* for FLW in the HvDRR33 sub-population by comparing RIL Hv.2018S.7.03209 and Hv.2018S.7.03211 carrying Georgie allele, and Hv.2018S.7.03369 carrying Lakhan allele at this QTL. A significantly narrower FL and L2 was observed for the RILs with Georgie allele compared to the one with Lakhan allele at *qHvDRR-FLS-17*. This trend was in accordance with the QTL effect observed in the QTL analysis and also in line with the parental inbreds, where Georgie had narrower leaves than Lakhan (Fig. 7a).

**Fig. 7.**
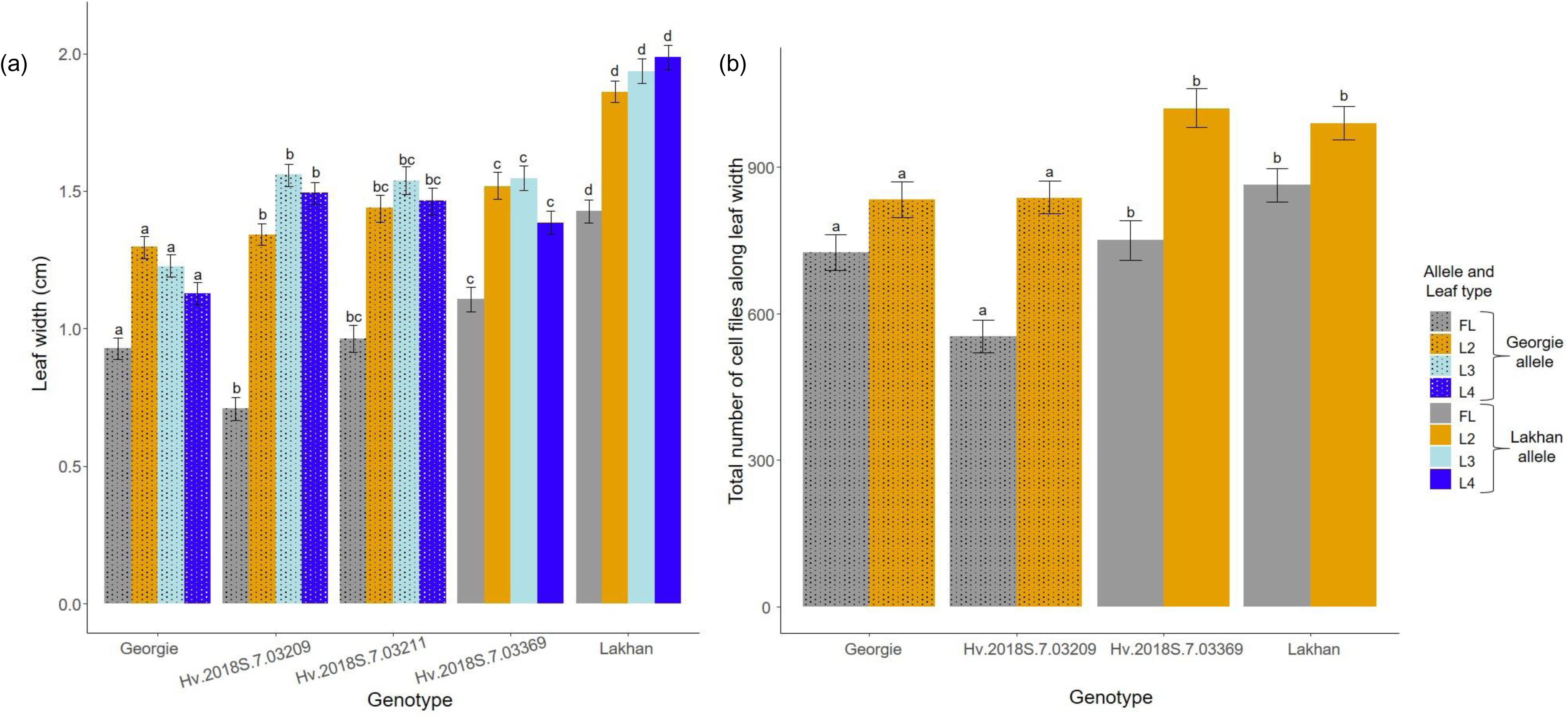
Validation of a major QTL *qHvDRR-FLS-17* associated with flag leaf length. *qHvDRR-FLS-17* was detected on chr4H and the validation was done in the HvDRR33 sub-population. (a) The width of flag leaf and older leaves in RILs and parental inbreds of sub-population HvDRR33. RILs were polymorphic for *qHvDRR-FLS-17* but isogenic for the other QTLs in the HvDRR33 sub-population. (b) The number of cell files along the leaf width. Bar indicates leaf width/total number of cell files along the leaf width (AEM ±SE, n = 6). Grey, yellow, light blue, and dark blue bar represents flag leaf (FL), second (L2), third (L3), and fourth leaf from the top (L4), respectively. Patterned bars in the graph represent Georgie or RILs with Georgie allele and non-patterned bars represent Lakhan or RILs with Lakhan allele at *qHvDRR-FLS-17*. Bars of the same color not sharing the same letter above the bar are significantly different from each other (*p* < 0.05). AEM, adjusted entry mean; SE, standard error.

Next, we estimated the total number of cells files across the leaf width and the average width of lateral cell (Fig. 7b, S9). At the QTL *qHvDRR-FLS-17*, the difference in FLW between the RIL pairs was explained by the total number of cell files along the leaf width (Fig. 7b).

### Fine mapping of QTL *qHvDRR-FLS-8* on chr2H

For fine mapping, we obtained the segregating population from the genetic cross between the above mentioned RILs from the HvDRR14 sub-population, Hv.2018S.7.02932 and Hv.2018S.7.02921 that are carrying Georgie allele and HOR8160 allele, respectively at *qHvDRR-FLS-8*. To enhance the resolution, 222 F2 plants that were recombinant between the markers flanking the QTL were further genotyped inside the interval. Later, F3 seeds of those recombinants were grown in the field and FLL was evaluated at the heading stage and the FLL ranged from 4.8 – 18.2 cm in these recombinants in the field. The *H^2^* was 48% for FLL in this segregating population. Collectively, the genotypic data of F2 population and phenotypic data from F3 population was used for single marker regression analysis. The segregating non-recombinants carrying Georgie allele at all the markers had significantly longer flag leaves than the individuals carrying HOR8160 allele which validated the QTL additionally (Fig. 8). The heterozygous individuals in the QTL region mimicked the phenotype of the individuals carrying the HOR8160 allele. The recombination analysis allowed us to delimit the QTL interval between M7 to M47 (626.5 to 629.9 Mb) consisting of 77 high confidence genes inside the new interval.

**Fig. 8.**
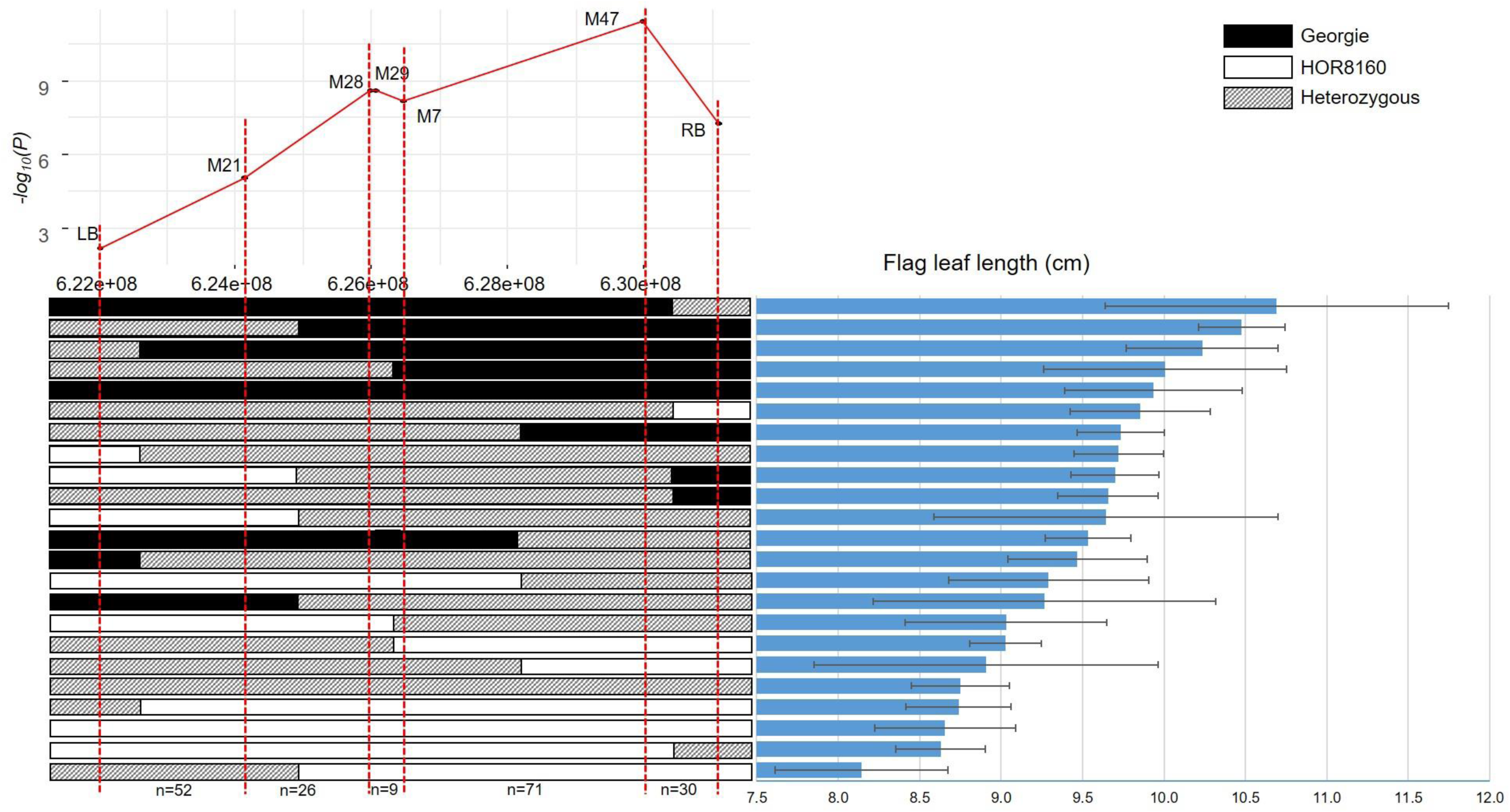
Fine mapping of *qHvDRR-FLS-8* on chromosome 2H. The figure on the left is the visual representation of the genotype of the individuals at the corresponding markers in the QTL interval. The black, white and grey bar represents Georgie, HOR8160 and heterozygous allele, respectively. The number below the left figure represents the number of recombinants between the respective adjacent markers. The graph on the right is the flag leaf length (AEM ±SE) of the corresponding individual. The top graph is the -log_10_(*p*) of single marker regression analysis performed using the genotypic data of F2 individuals and phenotypic data of F3 individuals.

## Discussion

Leaf size is unquestionably a key agronomic phenotype as the leaf is the main source of photosynthetic assimilates. Leaf size differences, and flag leaf in particular, are associated with yield potential in different plant species (Williams and Hayes, 1979). With this study, we aimed to understand the genetic complexity of FLL and FLW variation in barley.

### Flag leaf size variation and its association with the geographic origin

The length and width of the flag leaf of inbreds from warm regions of the world was bigger than those from cold regions (Fig. 1). The observation contradicts the theory of day time energy budget (carbon assimilation vs water loss) where bigger leaves are disadvantageous in warmer regions (Wright et al., 2017). One explanation for our finding might be that spring barley is cultivated from autumn (sown in Oct/Nov) to spring season (harvested in March/April) in warm and arid regions (Tabarzad et al., 2016), which reduces the high temperature constraint. On the other hand, spring barley in Europe and North America is grown from spring to summer (Wiersma and Ransom, 2005, Marcinkowski and Piniewski, 2018), which rules out that smaller leaves compensate for frost tolerance. However, the aforementioned trend that inbreds from warm regions had longer and wider flag leaves than those from colder regions was evident even within the European continent. FLL and FLW were bigger in the inbreds from Southern Europe than those from Northern Europe. Likewise, the inbreds from Central Asia with harsh winter showed FLL and FLW characteristics similar to those of Europe and North-America indicating a consistent spread of flag leaf characters along the temperature gradient.

It is also possible that the difference in leaf size can be compensated through the number of tillers and leaf number per culm in the inbreds from the temperate regions. Such studies at the population level are not available. However, on a physiological level, barley plants produce more tillers and final shoot biomass when grown at 15°C compared to those at 25°C (Hemming et al., 2012). Hence, the FLS alone might not be a good predictor of vegetation model for energy balance theory in barley and grasses in general. One approach to better understand this leaf size and temperature relationship on an ecological scale is to assess the effect of the individual genes contributing to leaf size variation.

### Genetic factors play major role in controlling flag leaf size characteristics in barley

In our study, we identified a high genetic contribution in defining FLS characteristics in HvDRR, with the broad-sense heritability (*H^2^*) on an entry mean basis of around 77% for FLL and around 58% for FLW (Table 1). The *H^2^* of FLL and FLW observed in our study corresponded to previous reports (Liu et al., 2015, Digel et al., 2016, Alqudah et al., 2018, Jabbari et al., 2018, Du et al., 2019) indicating a significant component of natural variation is under genetic control enabling isolation of causative genes.

The range of FLL (5.6 – 19 cm) and FLW (0.25 to 2.03 cm) observed in RILs of the HvDRR population was greater than those reported by Jabbari et al. (2018) in an association mapping panel indicating greater phenotypic diversity in the HvDRR population. This was further supported by the observation that phenotypic variation for FLL and FLW among the RILs of HvDRR was similar to those observed in the diversity panel evaluated in the current study. This finding illustrated the potential of the HvDRR population for genetic, physiological and ecological analyses.

The ranges reported by bi-parental population studies for FLL can be divided into two groups, the first one that observed wide range of phenotypic variation: Gyenis et al. (2007) recorded range of 8-34 cm; Vafadar Shamasbi et al. (2017), 3.9-13.4 cm; Niu et al. (2022), 5-31 cm. The high range of variation can be attributed to favorable controlled conditions (Liu et al., 2015, Digel et al., 2016, Alqudah et al., 2018) or single selected environmental conditions (Du et al., 2019) that were used in the respective studies. On the other side, a limited range of variation for FLL was reported for some studies, 17-19 cm (Liu et al., 2015), 12-14 cm (Du et al., 2019). In addition, the phenotypic range of FLW in previous studies in barley (Gyenis et al., 2007, Liu et al., 2015, Vafadar Shamasbi et al., 2017) was also narrower than those observed in our study. The above described limited range of FLL and FLW can be explained by the fact that those studies were conducted in individual bi-parental populations and the genetic divergence among the two parental genotypes was low. In contrast, the parents of the HvDRR population were selected to maximize genotypic and phenotypic diversity (Weisweiler et al., 2019).

### Genetic complexity of flag leaf size variation

Seventy-nine sub-population specific QTLs detected in our study coincided with those detected using MPP QTL analysis for flag leaf size variation in the current study (Fig. 3, 4). The significance of QTLs detected in the MPP analysis was higher than those detected in sub-population specific QTL analysis (Fig. 3, Table S3 and S4). We summarized these QTLs into 24 consensus QTLs. Furthermore, we compared the physical interval of the above mentioned 24 consensus QTLs with those reported in previous studies, whenever possible. We found that seven of the 24 consensus QTLs have been reported previously, which highlights the accuracy of the QTLs detected in our study (Table S5). On the other side, however, we detected 17 novel loci that explained between 3 to 44% phenotypic variance (Table S2, S3). This observation illustrated the potential and statistical power of a large and diverse multi-parent population to detect novel and rare variants. Nevertheless, our findings suggest that even all consensus QTL together explained just about 60% of the total phenotypic variance illustrating the genetic complexity of the studied traits.

Regardless of the significant genetic complexity of the investigated characteristics across all HvDRR populations, the proportion of the QTLs explaining 10% or more PEV was high. In addition, we observed an allelic series for the detected QTLs (Fig. 5). These findings suggest that fine mapping and eventually, reverse genetics approaches to isolate the underlying genes, will be possible for a substantial number of the detected QTLs when the most relevant sub-populations are studied.

### The function of some developmental genes might be conserved in crops

We detected QTLs for FLL and FLW harboring/residing close to known flowering time genes such as *Ppd-H1*, *Vrn-H1* and *Vrn-H2* (Table S5). In addition, 83% of the QTLs detected in our study co-localized with QTLs previously detected for grain size, flowering time, and plant height in the HvDRR population (Shrestha et al., 2022, Cosenza et al., 2024). Flowering time genes are reported to have pleiotropic effects on root and shoot growth (Arifuzzaman et al., 2016, Voss-Fels et al., 2018, Cosenza et al., 2024). For example, the late flowering allele of *Ppd-H1* (reduced photoperiod sensitivity) produced bigger leaves in barley (Alqudah et al., 2018) than the early ones (*Ppd-H1*, photoperiod sensitive). This report is in accordance to our observation that in the sub-populations HvDRR18 and HvDRR30, the parental inbred that contributed to early flowering allele at *Ppd-H1* locus had a smaller flag leaf than the ones associated with the late flowering allele. However, the opposite trend was observed for HvDRR26 and HvDRR39 where the early flowering allele at *Ppd-H1* was associated with longer and wider leaf (Table S8). This result contradicted those of Digel et al. (2016) where under long-day conditions *Ppd-H1* (late allele) produced bigger leaves then the genotypes carrying the early alleles. This finding might indicate that there is an additional gene located next to *Ppd-H1* which contributes to leaf size variation in the sub-populations HvDRR26 and HvDRR39.

*Vrn-H1 and Vrn-H2* separates winter and spring type in barley. Although our parental inbreds have been classified as spring type (Cosenza et al., 2024), we detected heading date (Cosenza et al., 2024) and leaf size QTLs at *Vrn-H1 and Vrn-H2*. However, we have observed no association between the presence of the late flowering allele and bigger leaf size at QTLs colocalizing with *Vrn-H2* in HvDRR30, HvDRR33, and HvDRR43 (Table S9). Therefore, there might be a gene independent of *Vrn-H2* in aforementioned QTL interval which is regulating the leaf size. On the other hand, we reported for the first time that allelic variation of *Vrn-H1* influenced FLL in spring barley in long-day conditions. The 11 Kb long intron 1 regulates the cold responsiveness and the deletion mutation in intron 1 bypass the vernalization requirement in spring barley (Hemming et al., 2009). We found that an allelic series for the deletion mutation in intron 1 (Fig. S7) affecting the flowering time in spring barley (Table 2). The early flowering allele *Vrn-H1-3* was associated with longer flag leaf (Table 2). In addition to the intron deletion, we found that hyper methylation of 116 bp region in intron 1 (chr5H: 528153135-528153251) (Kühl et al., 2025) reduced flowering time and increased FLL. Hence, our study uncovered previously unknown association of *Vrn-H1* with flag leaf size in spring barley.

The pleiotropic effect of the flowering time loci on above-ground biomass can be attributed to the duration of vegetative growth before transition to the reproductive phase (Neumann et al., 2017). However, the widely accepted correlation between early flowering and smaller leaf did not align perfectly with the flowering time associated loci in our study. This might be caused by the differences in the number of leaves or tillers between early and late flowering genotypes compensating for the differences in leaf size.

In addition to flowering time genes, we also detected QTLs that harbored orthologs of genes involved in leaf size variation in other plant species (Table S5). For instance, in the consensus QTL *qHvDRR-FLS-2* resides a barley gene (*HORVU.MOREX.r3.1HG0077830*) which is an ortholog of *Tcp5*. *Tcp5* belongs to the TCP family transcription factor containing protein and is the most promising candidate gene underlying *qHvDRR-FLS-2*. Yu et al. (2021) observed that *TCP5*-*KNAT3/SAW1* pathway plays an important role in defining leaf margin in Arabidopsis.

The consensus QTL *qHvDRR-FLS-15* on chr4H in our study overlapped with the previously detected QTL cluster C4 (Du et al., 2019) and this region harbors the barley ortholog (*HORVU.MOREX.r3.4HG0339430*) of Arabidopsis Growth-Regulating Factor (*GRF5*) (Table S5). *Grf5* belongs to the plant specific family of transcription factors which are known to be involved in different plant development processes (Hewezi et al., 2012, Kim et al., 2012, Liang et al., 2013, Bao et al., 2014, Debernardi et al., 2014, Liu et al., 2014, Pajoro et al., 2014). In Arabidopsis, *GRF5* was seen to interact with *AN3* and manipulate cell proliferation in leaf primordia (Horiguchi et al., 2005). The polymorphisms in *Tcp5* and *Grf5* genes between the small and large FL were located in the promoter region (Fig. S5 and S6) indicating a possible *cis*-regulatory differences.

Also, we found *HORVU.MOREX.r3.4HG0402190*, which is the ortholog of *NARROW LEAF7* (*Nal7*) residing in the consensus QTL *qHvDRR-FLS-16* on chr4H. *Nal7* belongs to Flavin-containing monoxygenase family and known to control the width of leaf blade in rice (Fujino et al., 2008). Hence, the function of barley ortholog of *Tcp5, Nal7* and *Grf*5 might be conserved and worth investigating in barley using transgenesis.

For other QTLs that explained high proportion of genotypic variance for leaf size, we didn’t find plausible candidate genes inside the confidence interval. The number of annotated genes in the confidence intervals, however, was too high, for a direct transgenic validation. Therefore, fine-mapping is one of the promising approaches to isolate underlying genes for such QTLs.

In the present study, we fine-mapped QTL *qHvDRR-FLS-8* which is significantly associated with FLL. Using only eight markers across the initial confidence interval of 8.7 Mb, we were able to reduce the region by two fifth. Seventy seven annotated high confidence genes are located in the reduced interval. Among them, 75 are polymorphic between the parental inbreds which makes a prioritization difficult. However, the number of recombination events between M7 and M47 is four, which will allow us to further narrow down the interval by genotyping additional markers in the interval (Fig. 8). Furthermore, additional ∼300 recombinants are selected in F4 generation.

### QTL *qHvDRR-FLS-8 and qHvDRR-FLS-17* also determine the size of leaves older than flag leaf

We characterized the effect of *qHvDRR-FLS-8* and *qHvDRR-FLS-17* in detail, as well as examined the anatomy of leaf epidermis to understand the link between leaf size and pavement cell size. Although these QTLs were detected based on the dimensions of FL, they also influenced the size of older leaves (Fig. 6, 7). For example, we saw that the leaf length was significantly different for the top three leaves, and progressively no vividness was observed for the fourth leaf. For leaf width, the significant differences were limited to the top two leaves.

The above findings are in contrast to results of the genes *Blf1* and *Nld1* which are known to influence the width of all the leaves in barley (Jöst et al., 2016, Yoshikawa et al., 2016). The same observation was made in rice for *Nal1* and *Nal2* (Ishiwata et al., 2013). Our observation is in agreement to results of Zanella et al. (2023) who studied a multi-parent population of wheat. In this study, *FLL5A* was observed to control the size of flag leaf but further the effect was gradually decreasing down towards the third leaf. One of the reasons that the behavior of QTLs detected in our study and that of Zanella et al. (2023) is different to the known genes influencing leaf size might be that the QTL studied here are representing the natural variation and not knock-out/overexpression lines. Therefore, the differences between alleles e.g. with respect to their gene expression but also protein functionality may be smaller.

It is known that leaf shape and size are the result of coordinated action of cell proliferation in primordium and cell expansion during the leaf blade elongation (Donnelly et al., 1999, Nath et al., 2003). For a better understanding of leaf size modulation, we characterized the QTLs *qHvDRR-FLS-8* and *qHvDRR-FLS-17* by analyzing the cells in the adaxial epidermal of the leaf blade. We measured the length of the lateral cells adjacent to the stomatal rows. We found that the differences in FLS between parental inbreds was associated with lateral cell expansion. A similar behavior was observed in the parental inbreds of wheat. Zanella et al. (2023) reported that cell size determined the leaf length at *FLL5A.* When we compared the RILs polymorphic only at *qHvDRR-FLS-8*, we found that the difference in FLL was caused by cell division but not longitudinal epidermal cell expansion.

Further, RILs carrying allele associated with wider leaf blade at *qHvDRR-FLS-17* showed differences in the number of cell files on the lateral plane in both FL and L2, rather than wider cells. This finding is in accordance with results of Jöst et al. (2016), (2024) and Yoshikawa et al. (2016) who observed that in their mutant-based studies in barley, the changes in the leaf width was due to change in the cell file number.

## Conclusion

The study identified previously unknown association of longer and wider flag leaf in genotypes adapted to warm region in barley natural population. In addition, we provided a comprehensive genetic control of FLL and FLW variation in barley and detected 17 novel genetic loci associated with flag leaf size. Furthermore, we identified sub-populations for fine-mapping major effect QTLs detected in the study. We also exemplified it by fine-mapping of *qHvDRR-FLS-8* and we are one step closer to isolate causal gene. Therewith, our study opens the door to functional validation of isolated candidate genes using transgenesis to unravel the underlying basis of the leaf size gradient across the globe but also optimize leaf size with respect to carbon assimilation.

## Acknowledgements

The authors would like to thank Konstantin Shek and Stephanie Krey (Quantitative Genetics and Genomics of Plants, Heinrich-Heine-University) for field and greenhouse management in Dusseldorf and Cologne. We are also grateful to Vasantha Nagam, Anna-Lena Hippel, Bjarne Michelsen, Uwe Gräf, Brigit Lemcke, Benjamin Brüdigam, Petra Müller, Grit Waldbauer, Jens Rokitte, Bernd Keding and Bernd Hackauf (Institute for Breeding Research on Agricultural Crops, Julius Kühn Institute) for their support in management of greenhouse and field experiments in Groß Lüsewitz. We also thank our student helpers Michael-Tobias Borghorst, Anabelle Odebrecht and Maximilian Maßling (Heinrich-Heine-University) for their help in leaf sampling. Finally, we would also like to thank Anjali Walpola, Aashu and Kefas Baiyi (Institute for Breeding Research on Agricultural Crops, Julius Kühn-Institute) for offering helping hands during field harvesting.

We would like to thank Deutsche Forschungsgemeinschaft (DFG, German Research Foundation) for financially supporting this work of the project “Fitness effect of molecular variants of leaf area and economics spectrum of Hordeum vulgare” within the transregional collaborative research center TRR341 ‘Plant Ecological Genetics’, TRR341. We also acknowledge the computational platform provided by the Centre of Information and Media Technology at Heinrich-Heine-University Düsseldorf and the department for Digitalisation and Artificial Intelligence of the Julius Kühn-Insitute.

## Author Contributions

BS and AS conceptualized the study. BS acquired the funding. BS and AS supervised the work. TL, YG, and AS conducted the experiments. TL, YG, PW, BS, and AS analyzed the data. DVI and AS designed and executed the field trials. TL, BS, and AS wrote the manuscript. TL and YG equally contributed to this work. All authors read, corrected, and approved the manuscript.

## References

R Core Team (2023): A Language and Environment for Statistical Computing_. R Foundation for Statistical Computing, Vienna, Austria. Thermo Fischer Scientific. QuantStudio 5 Real-Time PCR System.

Alqudah, A. M. & Schnurbusch, T. 2015. Barley Leaf Area and Leaf Growth Rates Are Maximized during the Pre-Anthesis Phase. Agronomy, 5, 107–129.

Alqudah, A. M., Youssef, H. M., Graner, A. & Schnurbusch, T. 2018. Natural variation and genetic make-up of leaf blade area in spring barley. Theoretical and Applied Genetics, 131, 873–886.

Arifuzzaman, M., Günal, S., Bungartz, A., Muzammil, S., P. Afsharyan, N., Léon, J. & Naz, A. A. 2016. Genetic Mapping Reveals Broader Role of Vrn-H3 Gene in Root and Shoot Development beyond Heading in Barley. PLOS One, 11, e0158718.

Bao, M., Bian, H., Zha, Y., Li, F., Sun, Y., Bai, B., Chen, Z., Wang, J., Zhu, M. & Han, N. 2014. miR396a-Mediated Basic Helix–Loop–Helix Transcription Factor bHLH74 Repression Acts as a Regulator for Root Growth in Arabidopsis Seedlings. Plant and Cell Physiology, 55, 1343–1353.

Bayer, M. M., Rapazote-Flores, P., Ganal, M., Hedley, P. E., Macaulay, M., Plieske, J., Ramsay, L., Russell, J., Shaw, P. D. & Thomas, W. 2017. Development and evaluation of a barley 50k iSelect SNP array. Frontiers in plant science, 8, 1792.

Blum, A. 1985. Photosynthesis and transpiration in leaves and ears of wheat and barley varieties. Journal of experimental botany, 36, 432–440.

Casale, F., Van Inghelandt, D., Weisweiler, M., Li, J. & Stich, B. 2022. Genomic prediction of the recombination rate variation in barley–A route to highly recombinogenic genotypes. Plant Biotechnology Journal, 20, 676–690.

Cosenza, F., Shrestha, A., VAN Inghelandt, D., Casale, F. A., Wu, P. Y., Weisweiler, M., Li, J., Wespel, F. & Stich, B. 2024. Genetic mapping reveals new loci and alleles for flowering time and plant height using the double round-robin population of barley. J Exp Bot, 75, 2385–2402.

Covarrubias-Pazaran, G. 2016. Genome-assisted prediction of quantitative traits using the R package sommer. PloS one, 11, e0156744.

Debernardi, J. M., Mecchia, M. A., Vercruyssen, L., Smaczniak, C., Kaufmann, K., Inze, D., Rodriguez, R. E. & Palatnik, J. F. 2014. Post-transcriptional control of GRF transcription factors by microRNA miR396 and GIF co-activator affects leaf size and longevity. The Plant Journal, 79, 413–426.

Digel, B., Tavakol, E., Verderio, G., Tondelli, A., Xu, X., Cattivelli, L., Rossini, L. & Von Korff, M. 2016. Photoperiod-H1 (Ppd-H1) controls leaf size. Plant Physiology, 172, 405–415.

Donald, C. M. 1968. The breeding of crop ideotypes. Euphytica, 17, 385-403.

Donnelly, P. M., Bonetta, D., Tsukaya, H., Dengler, R. E. & Dengler, N. G. 1999. Cell cycling and cell enlargement in developing leaves of Arabidopsis. Developmental biology, 215, 407–419.

Druka, A., Franckowiak, J., Lundqvist, U., Bonar, N., Alexander, J., Houston, K., Radovic, S., Shahinnia, F., Vendramin, V., Morgante, M., Stein, N. & Waugh, R. 2011. Genetic Dissection of Barley Morphology and Development Plant Physiology, 155, 617–627.

Du, B., Liu, L., Wang, Q., Sun, G., Ren, X., Li, C. & Sun, D. 2019. Identification of QTL underlying the leaf length and area of different leaves in barley. Sci Rep, 9, 4431.

Dupuis, J. & Siegmund, D. 1999. Statistical Methods for Mapping Quantitative Trait Loci From a Dense Set of Markers. Genetics, 151, 373–386.

Fan, Y., Tian, Z., Yan, Y., Hu, C., Abid, M., Jiang, D., Ma, C., Huang, Z. & Dai, T. 2017. Winter night-warming improves post-anthesis physiological activities and sink strength in relation to grain filling in winter wheat (Triticum aestivum L.). Frontiers in Plant Science, 8, 992.

Fujino, K., Matsuda, Y., Ozawa, K., Nishimura, T., Koshiba, T., Fraaije, M. W. & Sekiguchi, H. 2008. NARROW LEAF 7 controls leaf shape mediated by auxin in rice. Mol Genet Genomics, 279, 499–507.

Garin, V., Wimmer, V., Mezmouk, S., Malosetti, M. & Van Eeuwijk, F. 2017. How do the type of QTL effect and the form of the residual term influence QTL detection in multi-parent populations? A case study in the maize EU-NAM population. Theoretical and Applied Genetics, 130, 1753–1764.

Gautam, H., Fatma, M., Sehar, Z., Iqbal, N., Albaqami, M. & Khan, N. A. 2022. Exogenously-sourced ethylene positively modulates photosynthesis, carbohydrate metabolism, and antioxidant defense to enhance heat tolerance in rice. International Journal of Molecular Sciences, 23, 1031.

Gladun, I. 1993. Distribution of assimilates from the flag leaf of rice during the reproductive period of development. Russ. J. Plant. Physiol., 40, 215–219.

Gyenis, L., Yun, S., Smith, K., Steffenson, B., Bossolini, E., Sanguineti, M. C. & Muehlbauer, G. 2007. Genetic architecture of quantitative trait loci associated with morphological and agronomic trait differences in a wild by cultivated barley cross. Genome, 50, 714–723.

Haseneyer, G., Stracke, S., Paul, C., Einfeldt, C., Broda, A., Piepho, H. P., Graner, A. & Geiger, H. H. 2010. Population structure and phenotypic variation of a spring barley world collection set up for association studies. Plant Breeding, 129, 271–279.

Hemming, M. N., Fieg, S., JAMES Peacock, W., Dennis, E. S. & Trevaskis, B. 2009. Regions associated with repression of the barley (Hordeum vulgare) VERNALIZATION1 gene are not required for cold induction. Molecular Genetics and Genomics, 282, 107–117.

Hemming, M. N., Walford, S. A., Fieg, S., Dennis, E. S. & Trevaskis, B. 2012. Identification of high-temperature-responsive genes in cereals. Plant Physiol, 158, 1439–50.

Hewezi, T., Maier, T. R., Nettleton, D. & Baum, T. J. 2012. The Arabidopsis MicroRNA396-GRF1/GRF3 Regulatory Module Acts as a Developmental Regulator in the Reprogramming of Root Cells during Cyst Nematode Infection Plant Physiology, 159, 321-335.

Horiguchi, G., Kim, G. T. & Tsukaya, H. 2005. The transcription factor AtGRF5 and the transcription coactivator AN3 regulate cell proliferation in leaf primordia of Arabidopsis thaliana. Plant J, 43, 68–78.

Hu, J., Zhu, L., Zeng, D., Gao, Z., Guo, L., Fang, Y., Zhang, G., Dong, G., Yan, M. & Liu, J. 2010. Identification and characterization of NARROW AND ROLLED LEAF 1, a novel gene regulating leaf morphology and plant architecture in rice. Plant molecular biology, 73, 283–292.

Ishiwata, A., Ozawa, M., Nagasaki, H., Kato, M., Noda, Y., Yamaguchi, T., Nosaka, M., Shimizu-Sato, S., Nagasaki, A., Maekawa, M., Hirano, H. Y. & Sato, Y. 2013. Two WUSCHEL-related homeobox genes, narrow leaf2 and narrow leaf3, control leaf width in rice. Plant Cell Physiol, 54, 779–92.

Jabbari, M., Fakheri, B. A., Aghnoum, R., Mahdi Nezhad, N. & Ataei, R. 2018. GWAS analysis in spring barley (Hordeum vulgare L.) for morphological traits exposed to drought. PLOS One, 13, e0204952.

Jennings, P. R. 1964. Plant type as a rice breeding objective. Crop Sci., 4, 13–15.

Jiang, D., Fang, J., Lou, L., Zhao, J., Yuan, S., Yin, L., Sun, W., Peng, L., Guo, B. & Li, X. 2015. Characterization of a null allelic mutant of the rice NAL1 gene reveals its role in regulating cell division. PloS one, 10, e0118169.

Jöst, M., Hensel, G., Kappel, C., Druka, A., Sicard, A., Hohmann, U., Beier, S., Himmelbach, A., Waugh, R. & Kumlehn, J. 2016. The INDETERMINATE DOMAIN protein BROAD LEAF1 limits barley leaf width by restricting lateral proliferation. Current Biology, 26, 903–909.

Jost, M., Soltani, O., Kappel, C., Janiak, A., Chmielewska, B., Szurman-Zubrzycka, M., Mckim, S. M. & Lenhard, M. 2024. The gain-of-function mutation blf13 in the barley orthologue of the rice growth regulator NARROW LEAF1 is associated with increased leaf width. J Exp Bot, 75, 850–867.

Kim, J.-S., Mizoi, J., Kidokoro, S., Maruyama, K., Nakajima, J., Nakashima, K., Mitsuda, N., Takiguchi, Y., Ohme-Takagi, M., Kondou, Y., Yoshizumi, T., Matsui, M., Shinozaki, K. & Yamaguchi-Shinozaki, K. 2012. Arabidopsis GROWTH-REGULATING FACTOR7 Functions as a Transcriptional Repressor of Abscisic Acid– and Osmotic Stress–Responsive Genes, Including DREB2A The Plant Cell, 24, 3393-3405.

Kühl, M., Wu, P.-Y., Shrestha, A., Engelhorn, J., Hartwig, T. & Stich, B. 2025. Investigating the barley methylome, its variation and association with genomic, transcriptomic, and phenotypic variation. bioRxiv, 2024.10.21.619366.

Li, W. Q., Zhang, M. J., Gan, P. F., Qiao, L., Yang, S. Q., Miao, H., Wang, G. F., Zhang, M. M., Liu, W. T. & Li, H. F. 2017. CLD 1/SRL 1 modulates leaf rolling by affecting cell wall formation, epidermis integrity and water homeostasis in rice. The Plant Journal, 92, 904–923.

Liang, G., He, H., Li, Y., Wang, F. & Yu, D. 2013. Molecular Mechanism of microRNA396 Mediating Pistil Development in Arabidopsis Plant Physiology, 164, 249-258.

Liu, H., Guo, S., Xu, Y., Li, C., Zhang, Z., Zhang, D., Xu, S., Zhang, C. & Chong, K. 2014. OsmiR396d-Regulated OsGRFs Function in Floral Organogenesis in Rice through Binding to Their Targets OsJMJ706 and OsCR4. Plant Physiology, 165, 160–174.

Liu, L., Sun, G., Ren, X., Li, C. & Sun, D. 2015. Identification of QTL underlying physiological and morphological traits of flag leaf in barley. BMC genetics, 16, 1–10.

Long, S. P., Zhu, X. G., Naidu, S. L. & Ort, D. R. 2006. Can improvement in photosynthesis increase crop yields? Plant, cell & environment, 29, 315–330.

Marcinkowski, P. & Piniewski, M. 2018. Effect of climate change on sowing and harvest dates of spring barley and maize in Poland. International Agrophysics, 32.

Mascher, M., Wicker, T., Jenkins, J., Plott, C., Lux, T., Koh, C. S., Ens, J., Gundlach, H., Boston, L. B., Tulpová, Z., Holden, S., Hernández-Pinzón, I., Scholz, U., Mayer, K. F. X., Spannagl, M., Pozniak, C. J., Sharpe, A. G., Šimková, H., Moscou, M. J., Grimwood, J., Schmutz, J. & Stein, N. 2021. Long-read sequence assembly: a technical evaluation in barley. The Plant Cell, 33, 1888–1906.

Nath, U., Crawford, B. C., Carpenter, R. & Coen, E. 2003. Genetic control of surface curvature. Science, 299, 1404–1407.

Neuffer M. G., E. H. C., and S. R. Wessler, 1997. Mutants of Maize.. Cold Spring Harbor Laboratory Press, New York.

Neumann, K., Zhao, Y., Chu, J., Keilwagen, J., Reif, J. C., Kilian, B. & Graner, A. 2017. Genetic architecture and temporal patterns of biomass accumulation in spring barley revealed by image analysis. BMC Plant Biology, 17, 1–12.

Niu, Y., Chen, T., Zheng, Z., Zhao, C., Liu, C., Jia, J. & Zhou, M. 2022. A new major QTL for flag leaf thickness in barley (Hordeum vulgare L.). BMC Plant Biology, 22, 305.

Pajoro, A., Madrigal, P., Muiño, J. M., Matus, J. T., Jin, J., Mecchia, M. A., Debernardi, J. M., Palatnik, J. F., Balazadeh, S., Arif, M., Ó’maoiléidigh, D. S., Wellmer, F., Krajewski, P., Riechmann, J.-L., Angenent, G. C. & Kaufmann, K. 2014. Dynamics of chromatin accessibility and gene regulation by MADS-domain transcription factors in flower development. Genome Biology, 15, R41.

Piepho, H.-P. & Möhring, J. 2007. Computing Heritability and Selection Response From Unbalanced Plant Breeding Trials. Genetics, 177, 1881–1888.

Qi, J., Qian, Q., Bu, Q., Li, S., Chen, Q., Sun, J., Liang, W., Zhou, Y., Chu, C., Li, X., Ren, F., Palme, K., Zhao, B., Chen, J., Chen, M. & Li, C. 2008. Mutation of the Rice Narrow leaf1 Gene, Which Encodes a Novel Protein, Affects Vein Patterning and Polar Auxin Transport. Plant Physiology, 147, 1947–1959.

Rahman, M. A., Haque, M., Sikdar, B., Islam, M. A. & Matin, M. N. 2013. Correlation analysis of flag leaf with yield in several rice cultivars. Journal of Life and Earth Science, 8, 49–54.

Schneider, C. A., Rasband, W. S. & Eliceiri, K. W. 2012. NIH Image to ImageJ: 25 years of image analysis. Nature Methods, 9, 671–675.

Shrestha, A., Cosenza, F., Van Inghelandt, D., Wu, P. Y., Li, J., Casale, F. A., Weisweiler, M. & Stich, B. 2022. The double round-robin population unravels the genetic architecture of grain size in barley. J Exp Bot, 73, 7344–7361.

Stich, B. 2009. Comparison of mating designs for establishing nested association mapping populations in maize and Arabidopsis thaliana. Genetics, 183, 1525–1534.

Tabarzad, A., Ghaemi, A. A. & Zand-Parsa, S. 2016. Barley grain yield and protein content response to deficit irrigation and sowing dates in semi-arid region. Modern Applied Science, 10, 193–207.

Tang, X., Gong, R., Sun, W., Zhang, C. & Yu, S. 2018. Genetic dissection and validation of candidate genes for flag leaf size in rice (Oryza sativa L.). Theoretical and Applied Genetics, 131, 801–815.

Vafadar Shamasbi, F., Jamali, S. H., Sadeghzadeh, B. & Abdollahi Mandoulakani, B. 2017. Genetic mapping of quantitative trait loci for yield-affecting traits in a barley doubled haploid population derived from clipper× sahara 3771. Frontiers in Plant Science, 8, 688.

Van Es, S. W., Silveira, S. R., Rocha, D. I., Bimbo, A., Martinelli, A. P., Dornelas, M. C., Angenent, G. C. & Immink, R. G. 2018. Novel functions of the Arabidopsis transcription factor TCP 5 in petal development and ethylene biosynthesis. The Plant Journal, 94, 867–879.

Vanraden, P. M. 2008. Efficient methods to compute genomic predictions. Journal of dairy science, 91, 4414-4423.

Voss-Fels, K. P., Robinson, H., Mudge, S. R., Richard, C., Newman, S., Wittkop, B., Stahl, A., Friedt, W., Frisch, M. & Gabur, I. 2018. VERNALIZATION1 modulates root system architecture in wheat and barley. Molecular plant, 11, 226–229.

Wang, H., Hou, L., Wang, M. & Mao, P. 2016. Contribution of the pod wall to seed grain filling in alfalfa. Scientific Reports, 6, 26586.

Wang, J., Wang, T., Wang, Q., Tang, X., Ren, Y., Zheng, H., Liu, K., Yang, L., Jiang, H. & Li, Y. 2022. QTL mapping and candidate gene mining of flag leaf size traits in Japonica rice based on linkage mapping and genome-wide association study. Molecular Biology Reports, 1-9.

Wang, N., Wang, X., Qian, Y., Bai, D., Bao, Y., Zhao, X., Xu, P., Li, K., Li, J. & Li, K. 2023. Genome-Wide Association Analysis of Rice Leaf Traits. Agronomy, 13, 2687.

Weisweiler, M., Arlt, C., Wu, P. Y., Van Inghelandt, D., Hartwig, T. & Stich, B. 2022. Structural variants in the barley gene pool: precision and sensitivity to detect them using short-read sequencing and their association with gene expression and phenotypic variation. Theor Appl Genet, 135, 3511–3529.

Weisweiler, M., Montaigu, A., Ries, D., Pfeifer, M. & Stich, B. 2019. Transcriptomic and presence/absence variation in the barley genome assessed from multi-tissue mRNA sequencing and their power to predict phenotypic traits. BMC Genomics, 20, 787.

Wen, Y., Fang, Y., Hu, P., Tan, Y., Wang, Y., Hou, L., Deng, X., Wu, H., Zhu, L. & Zhu, L. 2020. Construction of a high-density genetic map based on SLAF markers and QTL analysis of leaf size in rice. Frontiers in Plant Science, 11, 1143.

Wiersma, J. J. & Ransom, J. K. 2005. The small grains field guide.

Williams, R. & Hayes, J. 1979. Relationships between photosynthetic area and other growth attributes with grain yield in 6- and 2-row barley genotypes. Annals of Applied Biology, 91, 391–395.

Wright, I. J., Dong, N., Maire, V., Prentice, I. C., Westoby, M., Díaz, S., Gallagher, R. V., Jacobs, B. F., Kooyman, R., Law, E. A., Leishman, M. R., Niinemets, Ü., Reich, P. B., Sack, L., Villar, R., Wang, H. & Wilf, P. 2017. Global climatic drivers of leaf size. Science, 357, 917–921.

Wu, J., Qi, Y., Hu, G., Li, J., Li, Z. & Zhang, H. 2017. Genetic architecture of flag leaf length and width in rice (Oryza sativa L.) revealed by association mapping. Genes & Genomics, 39, 341–352.

Wu, X. Y., Kuai, B. K., Jia, J. Z. & Jing, H. C. 2012. Regulation of leaf senescence and crop genetic improvement F. Journal of Integrative Plant Biology, 54, 936–952.

Xiang, J.-J., Zhang, G.-H., Qian, Q. & Xue, H.-W. 2012. Semi-rolled leaf1 encodes a putative glycosylphosphatidylinositol-anchored protein and modulates rice leaf rolling by regulating the formation of bulliform cells. Plant physiology, 159, 1488–1500.

Xue, D.-W., Chen, M.-C., Zhou, M.-X., Chen, S., Mao, Y. & Zhang, G.-P. 2008. QTL analysis of flag leaf in barley (Hordeum vulgare L.) for morphological traits and chlorophyll content. Journal of Zhejiang University Science B, 9, 938–943.

Yang, H., Fang, C., Li, Y., Wu, Y., Fransson, P., Rillig, M. C., Zhai, S., Xie, J., Tong, Z. & Zhang, Q. 2022. Temporal complementarity between roots and mycorrhizal fungi drives wheat nitrogen use efficiency. New Phytologist, 236, 1168–1181.

Yoshikawa, T., Tanaka, S.-Y., Masumoto, Y., Nobori, N., Ishii, H., Hibara, K.-I., Itoh, J.-I., Tanisaka, T. & Taketa, S. 2016. Barley NARROW LEAFED DWARF1 encoding a WUSCHEL-RELATED HOMEOBOX 3 (WOX3) regulates the marginal development of lateral organs. Breeding Science, 66, 416-424.

Yu, H., Zhang, L., Wang, W., Tian, P., Wang, W., Wang, K., Gao, Z., Liu, S., Zhang, Y., Irish, V. F. & Huang, T. 2021. TCP5 controls leaf margin development by regulating KNOX and BEL-like transcription factors in Arabidopsis. J Exp Bot, 72, 1809–1821.

Zanella, C. M., Rotondo, M., Mccormick-Barnes, C., Mellers, G., Corsi, B., Berry, S., Ciccone, G., Day, R., Faralli, M., Galle, A., Gardner, K. A., Jacobs, J., Ober, E. S., Sanchez Del Rio, A., VAN Rie, J., Lawson, T. & Cockram, J. 2023. Longer epidermal cells underlie a quantitative source of variation in wheat flag leaf size. New Phytol, 237, 1558–1573.

Zhang, B., Ye, W., Ren, D., Tian, P., Peng, Y., Gao, Y., Ruan, B., Wang, L., Zhang, G. & Guo, L. 2015. Genetic analysis of flag leaf size and candidate genes determination of a major QTL for flag leaf width in rice. Rice, 8, 1–10.

Zhang, G.-H., Xu, Q., Zhu, X.-D., Qian, Q. & Xue, H.-W. 2009. SHALLOT-LIKE1 Is a KANADI Transcription Factor That Modulates Rice Leaf Rolling by Regulating Leaf Abaxial Cell Development The Plant Cell, 21, 719-735.

Zheng, T. 1999. Effects of some photosynthetic organs on milking and grain yield of barley. Barley Science, 1, 21–22.

