## Supplementary Figures S1-S9 for "Genetic architecture and cellular basis of flag leaf size variation in barley"

### Slide 1
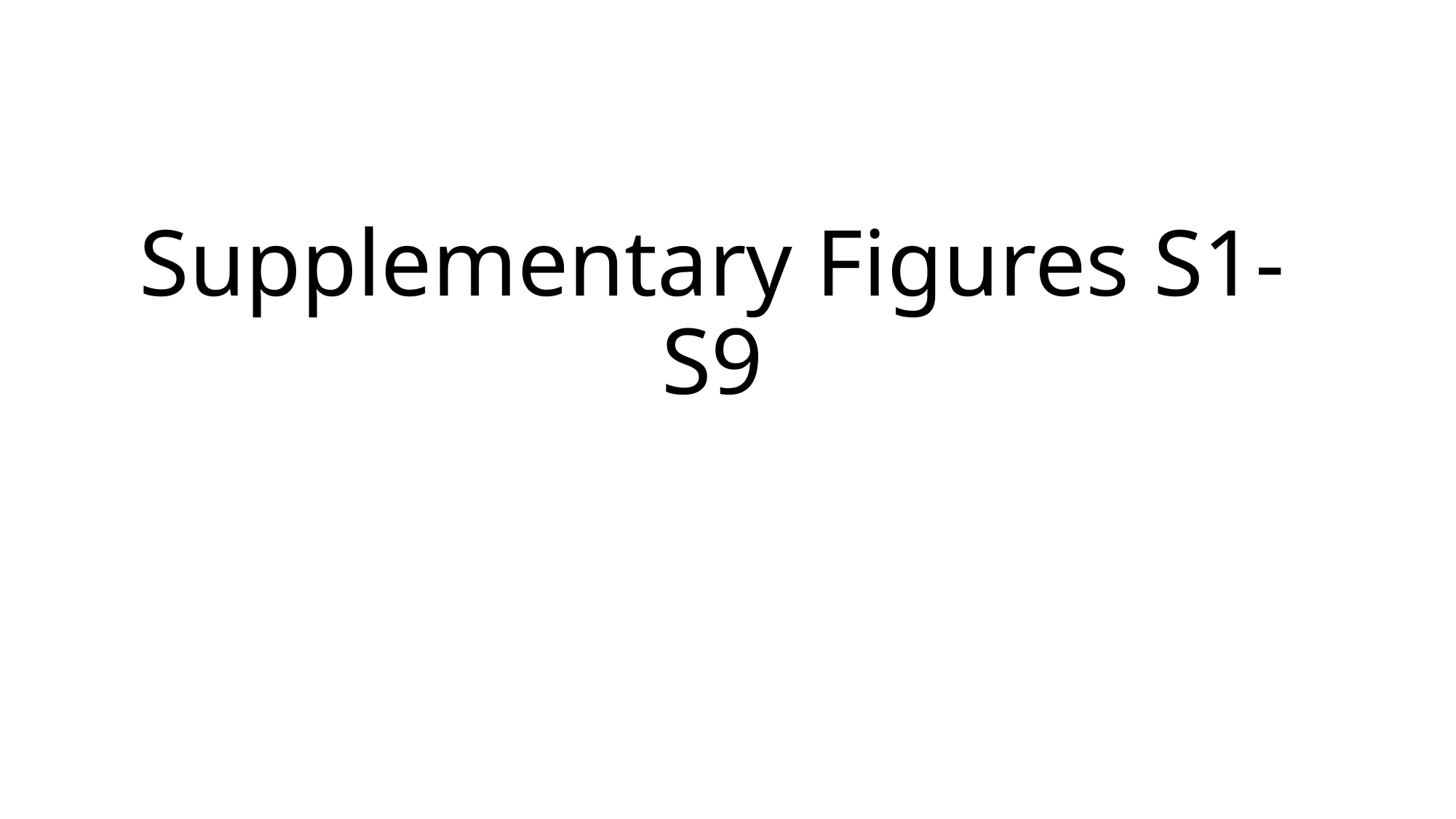

#### Supplementary Figures S1-S9

### Slide 2
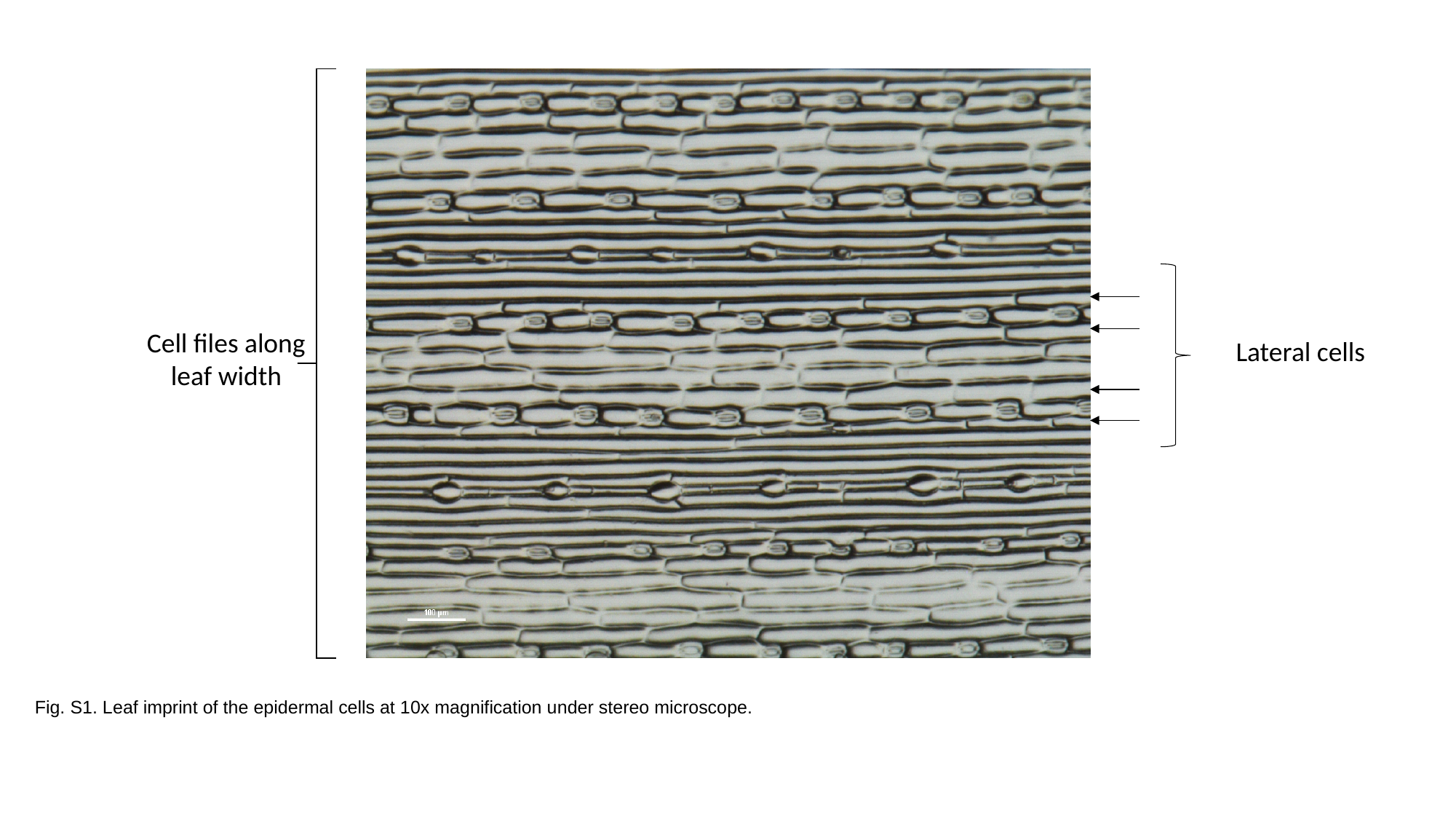

Cell files along leaf width
Lateral cells
Fig. S1. Leaf imprint of the epidermal cells at 10x magnification under stereo microscope.

### Slide 3
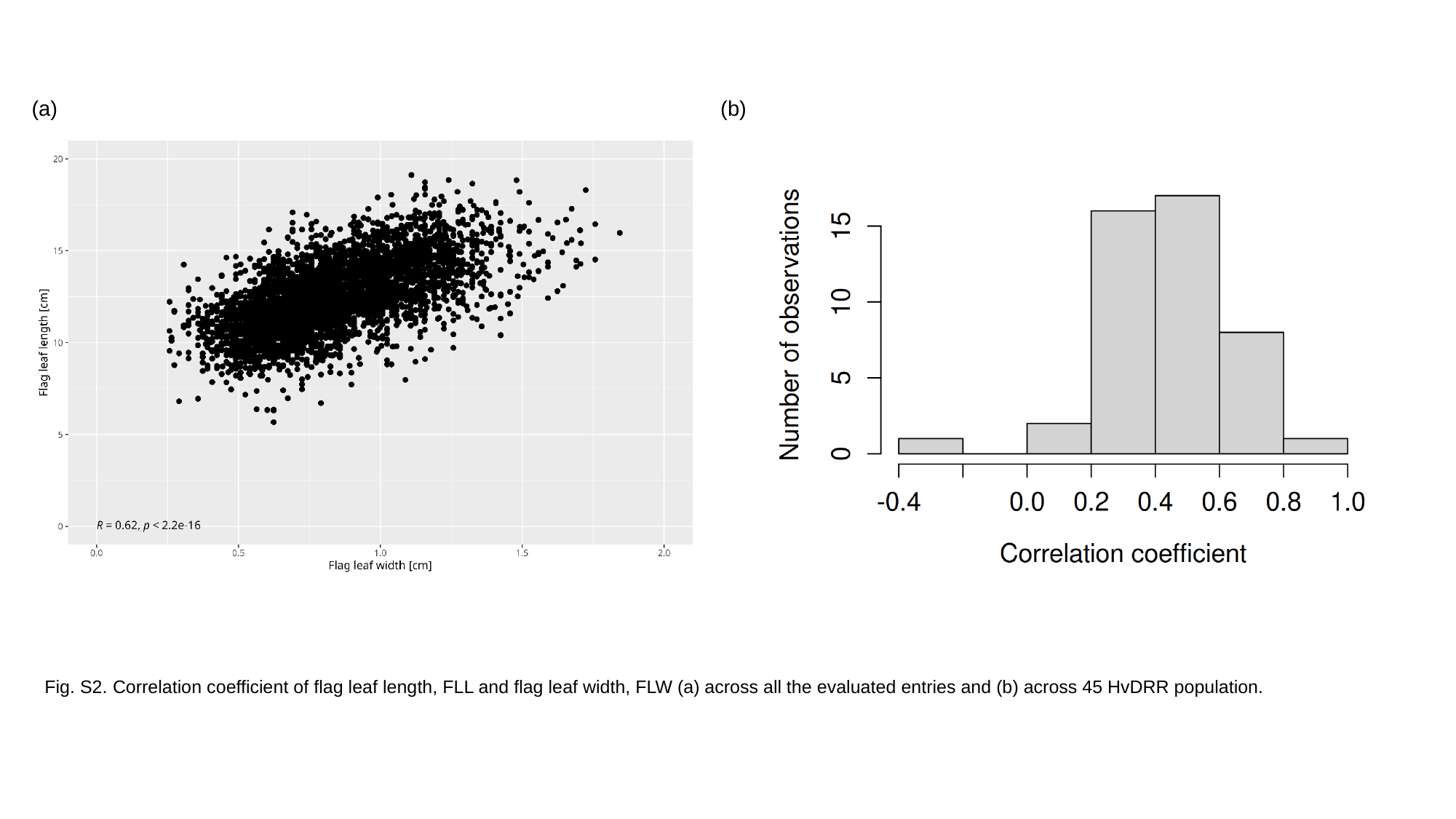

(b)
(a)
Fig. S2. Correlation coefficient of flag leaf length, FLL and flag leaf width, FLW (a) across all the evaluated entries and (b) across 45 HvDRR population.

### Slide 4
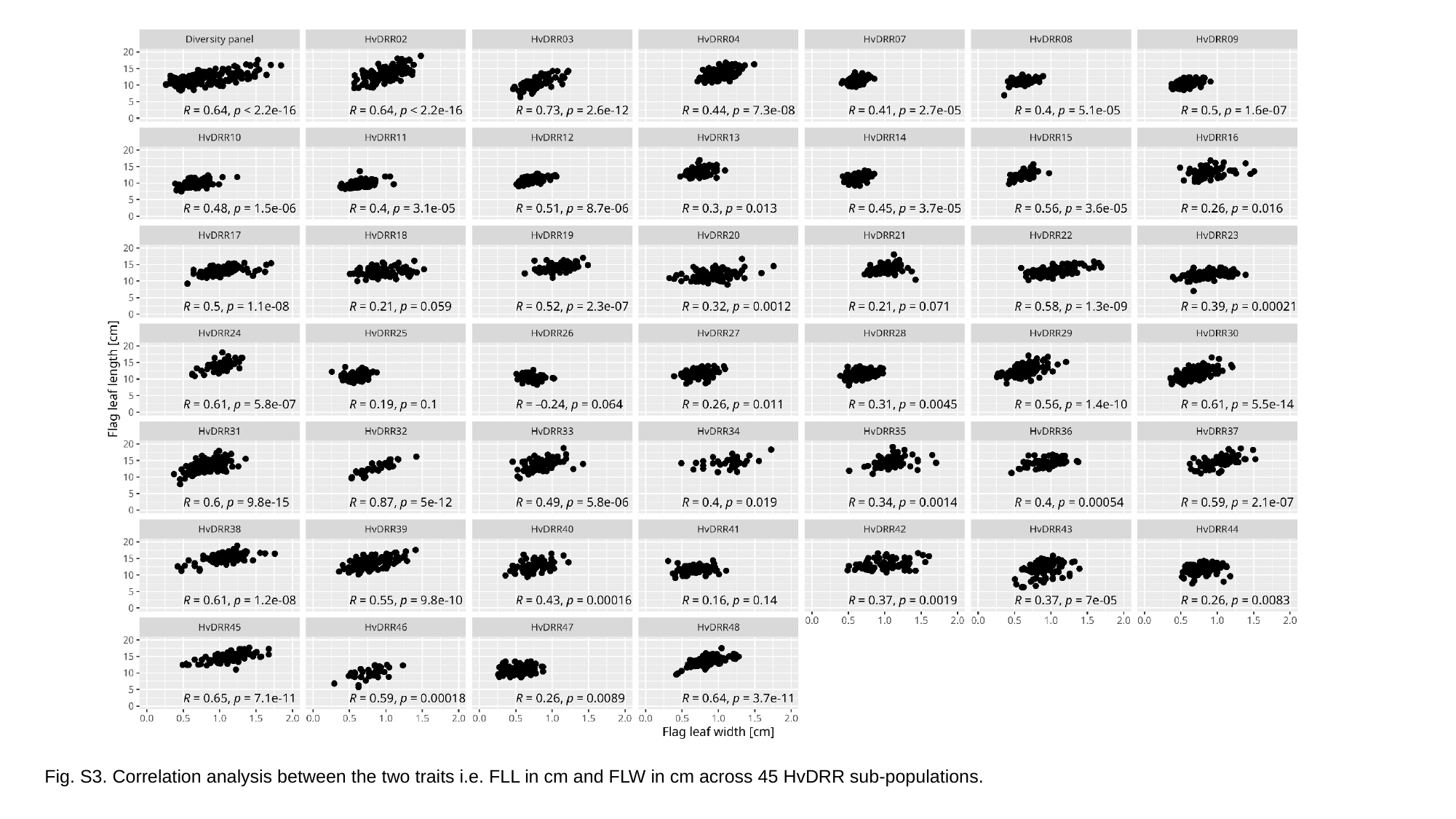

Fig. S3. Correlation analysis between the two traits i.e. FLL in cm and FLW in cm across 45 HvDRR sub-populations.

### Slide 5
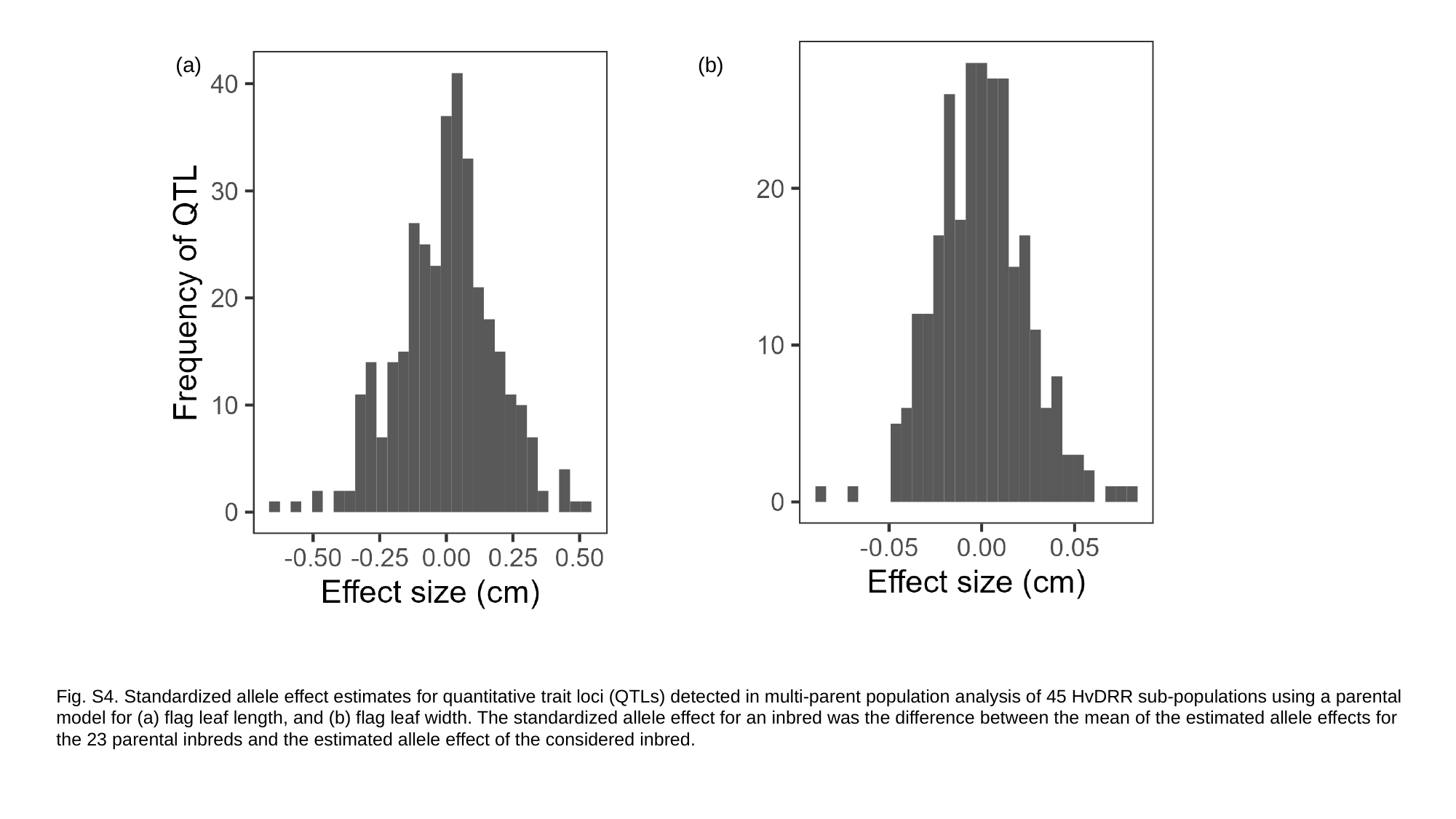

(a)
(b)
Fig. S4. Standardized allele effect estimates for quantitative trait loci (QTLs) detected in multi-parent population analysis of 45 HvDRR sub-populations using a parental model for (a) flag leaf length, and (b) flag leaf width. The standardized allele effect for an inbred was the difference between the mean of the estimated allele effects for the 23 parental inbreds and the estimated allele effect of the considered inbred.

### Slide 6
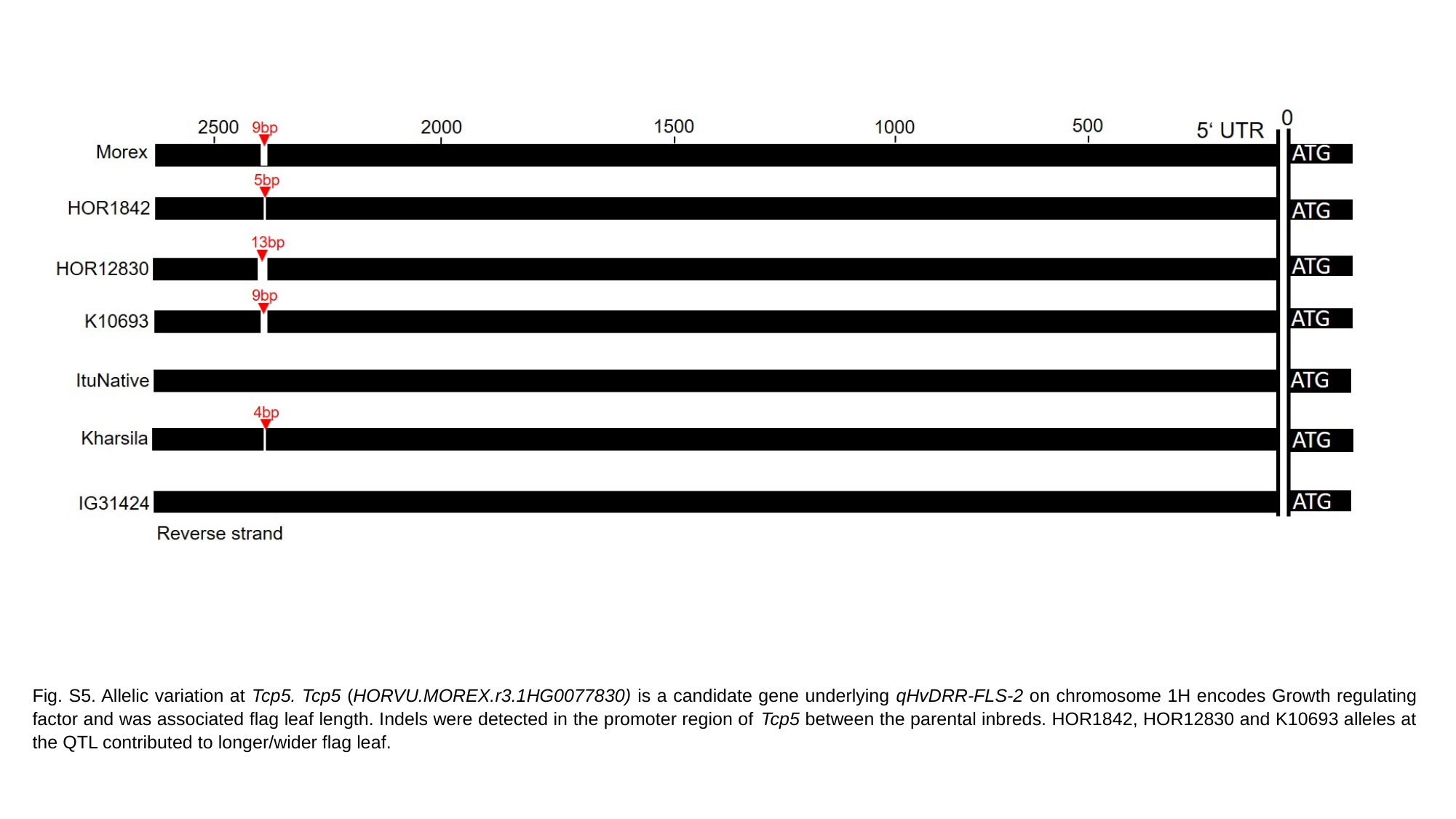

5‘ UTR
Fig. S5. Allelic variation at Tcp5. Tcp5 (HORVU.MOREX.r3.1HG0077830) is a candidate gene underlying qHvDRR-FLS-2 on chromosome 1H encodes Growth regulating factor and was associated flag leaf length. Indels were detected in the promoter region of Tcp5 between the parental inbreds. HOR1842, HOR12830 and K10693 alleles at the QTL contributed to longer/wider flag leaf.

### Slide 7
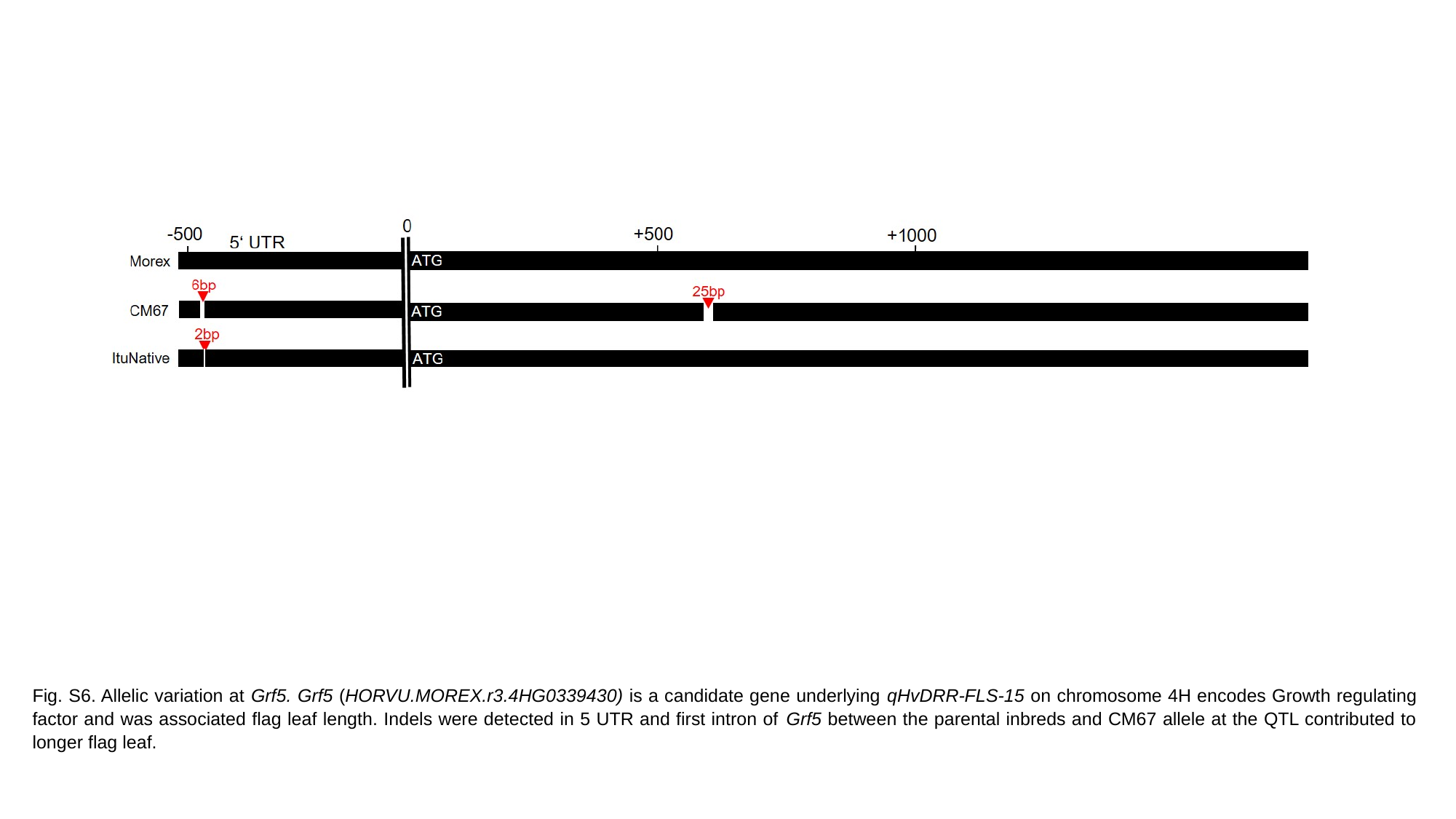

Fig. S6. Allelic variation at Grf5. Grf5 (HORVU.MOREX.r3.4HG0339430) is a candidate gene underlying qHvDRR-FLS-15 on chromosome 4H encodes Growth regulating factor and was associated flag leaf length. Indels were detected in 5 UTR and first intron of Grf5 between the parental inbreds and CM67 allele at the QTL contributed to longer flag leaf.

### Slide 8
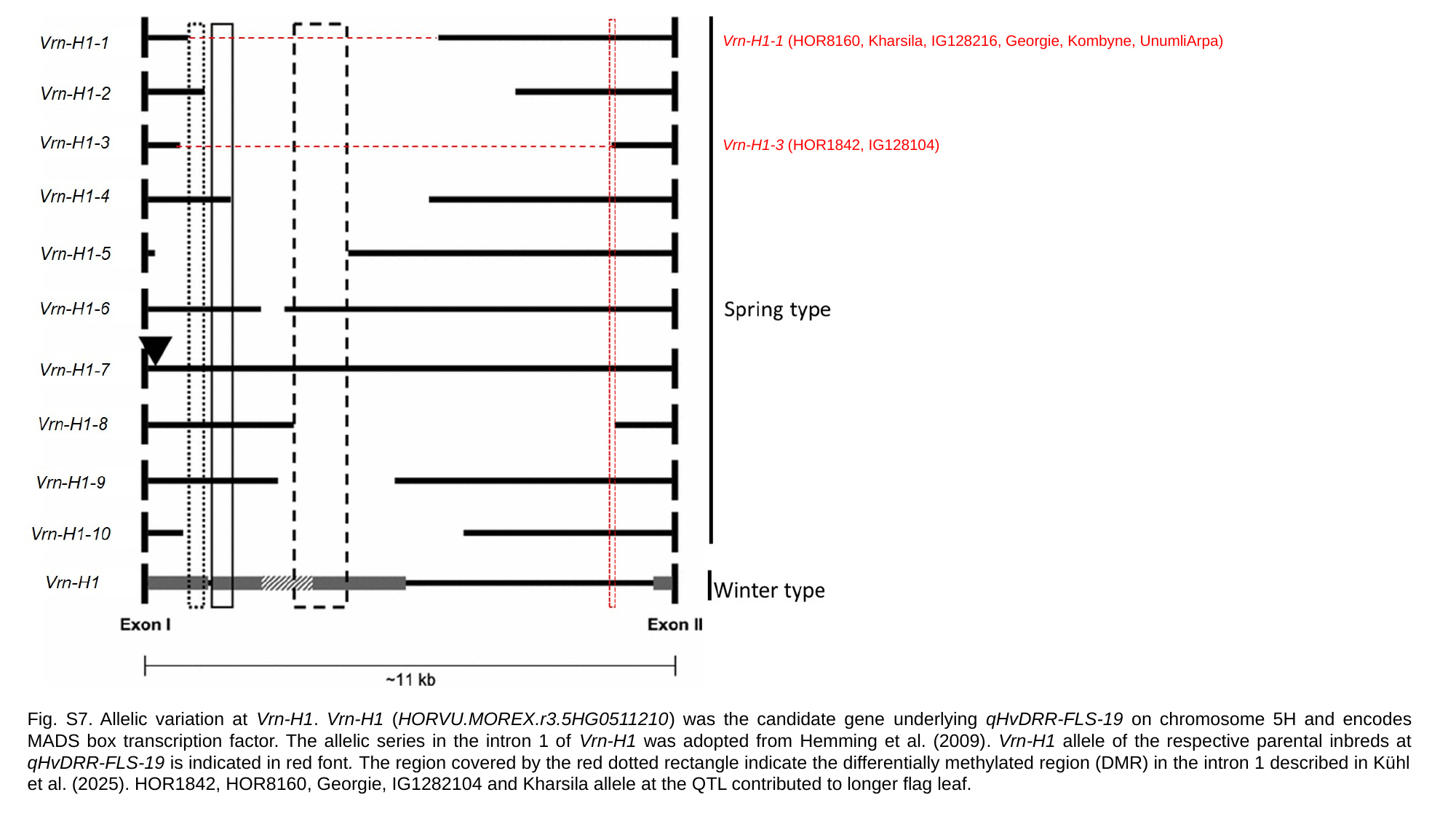

Vrn-H1-1 (HOR8160, Kharsila, IG128216, Georgie, Kombyne, UnumliArpa)
Vrn-H1-3 (HOR1842, IG128104)
Fig. S7. Allelic variation at Vrn-H1. Vrn-H1 (HORVU.MOREX.r3.5HG0511210) was the candidate gene underlying qHvDRR-FLS-19 on chromosome 5H and encodes MADS box transcription factor. The allelic series in the intron 1 of Vrn-H1 was adopted from Hemming et al. (2009). Vrn-H1 allele of the respective parental inbreds at qHvDRR-FLS-19 is indicated in red font. The region covered by the red dotted rectangle indicate the differentially methylated region (DMR) in the intron 1 described in Kühl et al. (2025). HOR1842, HOR8160, Georgie, IG1282104 and Kharsila allele at the QTL contributed to longer flag leaf.

### Slide 9
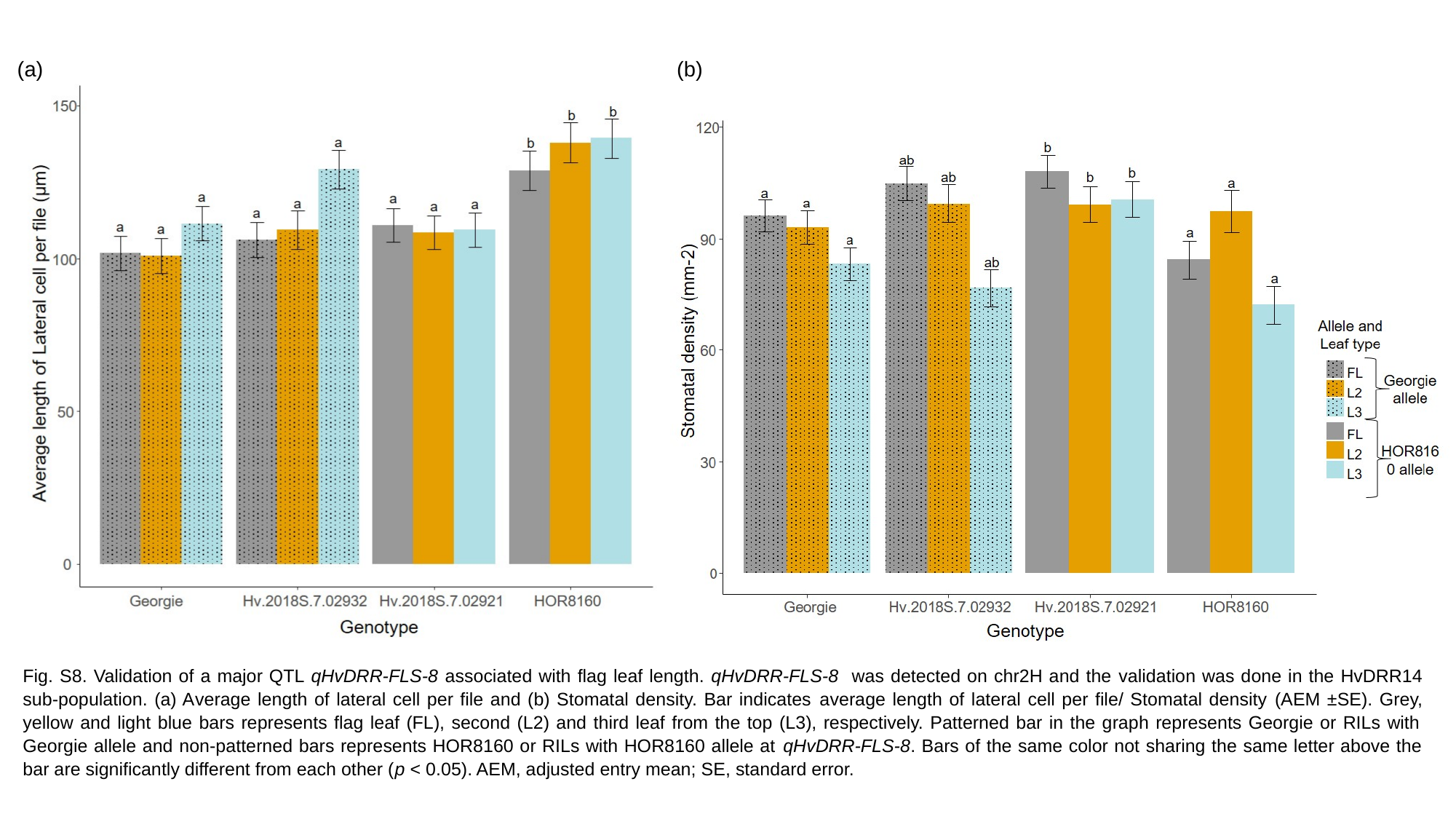

(b)
(a)
Fig. S8. Validation of a major QTL qHvDRR-FLS-8 associated with flag leaf length. qHvDRR-FLS-8 was detected on chr2H and the validation was done in the HvDRR14 sub-population. (a) Average length of lateral cell per file and (b) Stomatal density. Bar indicates average length of lateral cell per file/ Stomatal density (AEM ±SE). Grey, yellow and light blue bars represents flag leaf (FL), second (L2) and third leaf from the top (L3), respectively. Patterned bar in the graph represents Georgie or RILs with Georgie allele and non-patterned bars represents HOR8160 or RILs with HOR8160 allele at qHvDRR-FLS-8. Bars of the same color not sharing the same letter above the bar are significantly different from each other (p < 0.05). AEM, adjusted entry mean; SE, standard error.

### Slide 10
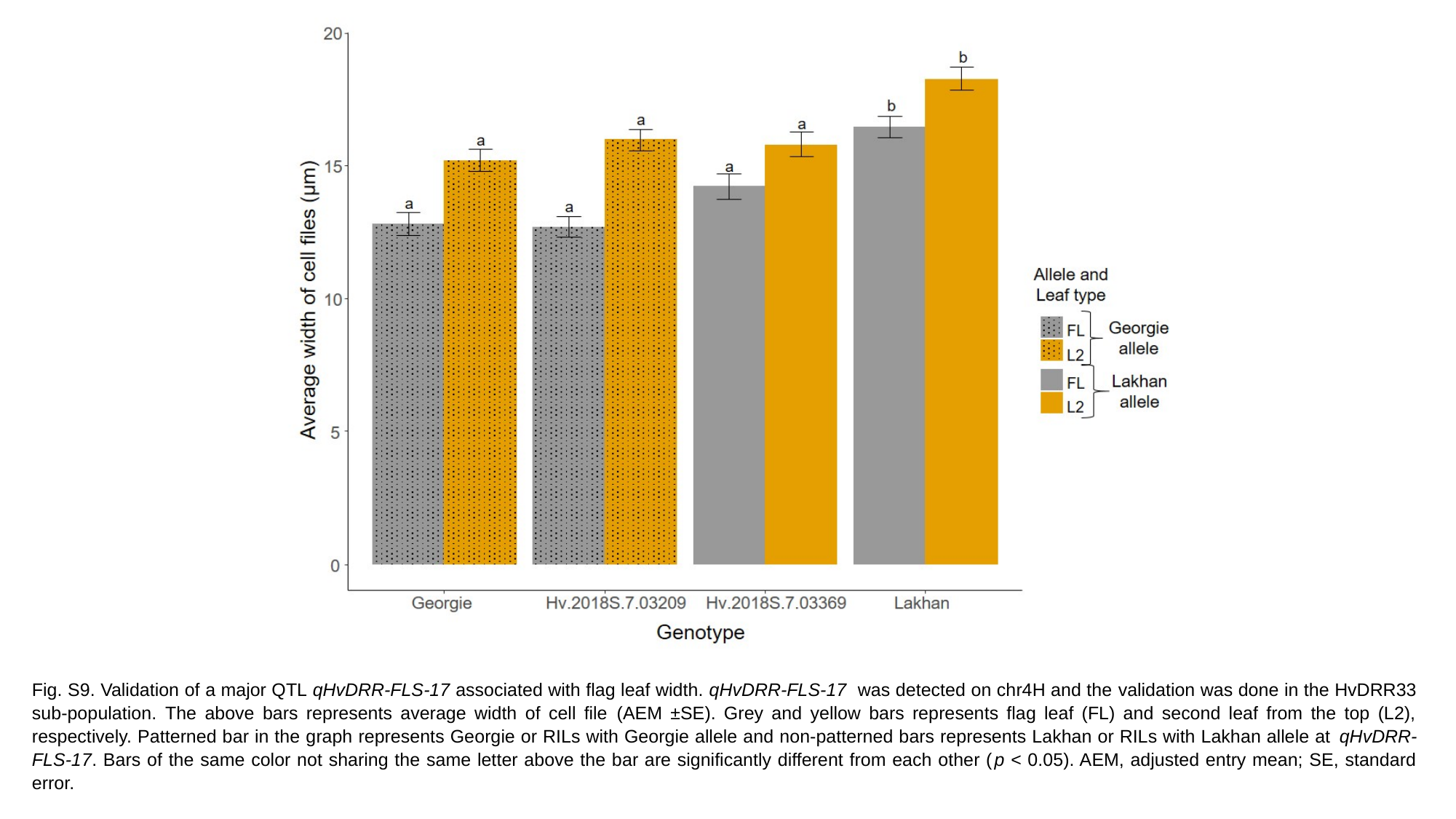

Fig. S9. Validation of a major QTL qHvDRR-FLS-17 associated with flag leaf width. qHvDRR-FLS-17 was detected on chr4H and the validation was done in the HvDRR33 sub-population. The above bars represents average width of cell file (AEM ±SE). Grey and yellow bars represents flag leaf (FL) and second leaf from the top (L2), respectively. Patterned bar in the graph represents Georgie or RILs with Georgie allele and non-patterned bars represents Lakhan or RILs with Lakhan allele at qHvDRR-FLS-17. Bars of the same color not sharing the same letter above the bar are significantly different from each other (p < 0.05). AEM, adjusted entry mean; SE, standard error.
